# The super-enhancer landscape reflects molecular subgroups of adrenocortical carcinoma

**DOI:** 10.1101/2023.04.05.535576

**Authors:** Samuel Gunz, Gwenneg Kerdivel, Jonas Meirer, Igor Shapiro, Bruno Ragazzon, Floriane Amrouche, Marie-Ange Calmejane, Juliette Hamroune, Sandra Sigala, Alfredo Berruti, Jérôme Bertherat, Guillaume Assié, Constanze Hantel, Valentina Boeva

## Abstract

Adrenocortical carcinoma (ACC) is a rare cancer of the adrenal gland with generally very unfavourable outcome. Two molecular subgroups, C1A and C1B, have been previously identified with a significant association with patient survival. In this work, we study chromatin state organization characterized by histone modifications using ChIP-sequencing in adult ACC. We describe the super-enhancer landscape of ACC, characterized by H3K27ac, and identify super-enhancer regulated genes that play a significant role in tumorigenesis. We show that the super-enhancer landscape reflects differences between the molecular sub-groups C1A and C1B and identify networks of master transcription factors mirroring these differences. Additionally, we study the effects of molecules THZ1 and JQ1 previously reported to affect super-enhancer-driven gene expression in ACC cell lines. Our results reveal that the landscape of histone modifications in ACC is linked to its molecular subgroups and thus provide the groundwork for future analysis of epigenetic reprogramming in ACC.

---

Adrenocortical carcinoma (ACC) is a rare yet aggressive endocrine cancer of the adrenal gland. Recently, comprehensive genomic characterization of ACC allowed to catalogue ACC driver genes and link other genomic features, such as certain somatic copy number alterations and whole genome doubling, with aggressive clinical course. Nevertheless, genetic profiles of ACC tumors are highly heterogeneous, with nearly 30% of the tumors lacking mutations in known driver genes [1, 2]. Transcriptomic studies revealed two molecular subgroups, C1A and C1B. C1A was associated with poor prognosis, while patients in the C1B subgroup showed significantly better outcomes [3, 4]. Most tumors of the aggressive C1A subgroup exhibited a steroid phenotype, which impacted prognosis [2].

Non-mutational epigenetic reprogramming is a hallmark of cancer that gained interest in cancer research in recent years [5]. In ACC, an analysis of DNA methylation identified a CpG Island Methylator Phenotype (CIMP) [1, 3]. The CIMP phenotype in ACC was linked to an especially poor prognosis, potentially due to the epigenetic silencing of tumor suppressor genes and genes involved in neoantigen presentation [6, 7]. In addition, epigenetic remodeling characterized by changes in histone post-translational modifications has been linked to tumorigenesis and the characterization of various cancer subgroups [8–11]. To date, little is known about chromatin regulation and reprogramming in ACC.

In this study, we investigate different broad chromatin domains and their presence in ACC. First, we show that superenhancers, regions of the genome where multiple enhancers are clustered together insuring quick and strong transcriptional response of target genes, regulate known ACC driver genes and genes whose expression is predictive of patient survival. Moreover, we uncover that the super-enhancer landscape reflects previously described molecular subgroups of ACC, C1A and C1B. We further characterize core regulatory circuitries (CRCs) of master transcriptional regulators in ACC and assess their importance in shaping the characteristics of the ACC molecular subgroups. Using ACC cell lines, we study the effects of BRD4 and CDK7 inhibitors, JQ1 and THZ1, reported to affect expression of super-enhancer-driven genes on the expression of CRC genes and on transcriptional programs linked to the ACC molecular subgroups. Second, we analyze super-silencers in ACC, *i*.*e*., large domains enriched in trimethylation of lysine 27 on histone H3 (H3K27me3) catalyzed by Polycomb Repressive Complex 2 (PRC2). We show that the super-silencers landscape discriminate cancer from non-cancer tissue in the adrenal gland. Third, we investigate the relationship between ACC molecular subgroups and trimethylation of lysine 4 on histone H3 (H3K4me3) domains marking active or poised promoters. Our results suggest that there may be a link between the variability in broad H3K4me3 domain patterns and the CIMP phenotype in ACC, indicating a need for future investigation.

## Results

### Characteristics of the super-enhancer landscape in ACC

Following the standard pipeline to identify super-enhancers, we generated and analyzed ChIP-seq data of the histone modification marking active enhancers and promoters, histone H3 lysine 27 acetylation (H3K27ac) [12], in 15 ACC tumor samples and 2 samples of non-cancer tissue of the adrenal gland (Figure 1a-b and Methods). We identified a total of 3742 super-enhancer regions to be active in at least two cancer or non-cancer samples. For each super-enhancer region, we calculated a score in all analyzed samples by summing the normalized H3K27ac ChIP-seq signal in the respective region. This score was further used to assign super-enhancers to their gene targets resulting in 2836 regions with gene associations (Methods).

**Figure 1:**
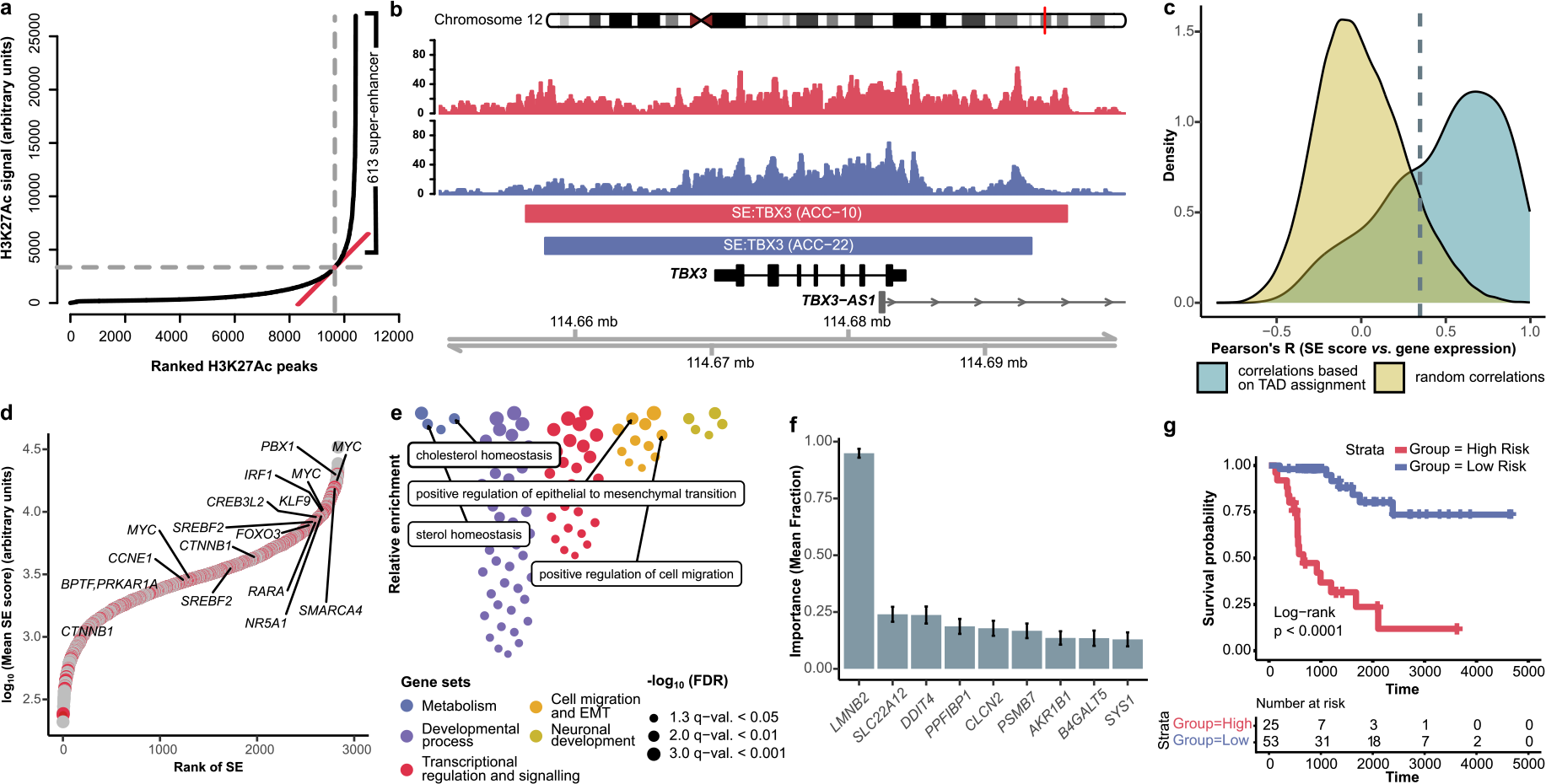
Characterization of the super-enhancer landscape of ACC. **a**, Visualization of the super-enhancer calling procedure based on the strength of the H3K27ac signal in ACC sample 22; H3K27ac peaks were stitched together and ranked according to their aggregated signal strength, the inflection point above which the peaks were classified as super-enhancers (SEs) was determined by the ROSE [12] algorithm. **b**, Example of a super-enhancer associated with the *TBX3* gene detected using the H3K27ac ChIP-seq signal; ACC samples 10 (red) and 22 (blue) are shown. **c**, Distribution of Pearsons’s correlation coefficients between super-enhancer scores and expression values of putative target genes showing strong association between activity of super-enhancers and gene expression; Correlations with expression of genes in the TAD neighborhood (blue) and random genes (yellow). The dashed line indicates the correlation threshold determined using the 90% quantile of the distribution for random associations. **d**, Super-enhancers ranked by the mean activity score over all ACC samples. Labeled are super-enhancer target genes that were previously described in relation to ACC. Red circles correspond to target genes representing transcription factors and/or known oncogenes. **e**, Gene ontology analysis of super-enhancer target genes. The colors indicate functional classes. Highlighted in boxes are GO terms relevant for the ACC pathogenesis. **f**, Variable importance of super-enhancer target genes used in survival analysis. The plot shows the fraction of time each of the 20 genes with highest predictive performance was selected in the regularized survival analysis. **g**, Kaplan–Meier overall survival curves for high- and low-risk patient groups determined by the expression values of the super-enhancer-regulated genes selected by our regularized survival model. SE, super-enhancer; FDR, false discovery rate.

Ranking the super-enhancer regions by their average activity score over all ACC samples revealed that super-enhancers with highest scores were largely associated with transcription factors (TFs) and/or oncogenes. Among them, we identified a number of genes previously described in the context of ACC, including *PBX1* [13, 14], *MYC* [15]), *SMARCA4*/*BRG1* [16, 17], *NR5A1*/*SF-1* [2, 14, 18], *ZNFR3* [1, 2], *CTNNB1* [1, 2], *CCNE1* [2], and *PRKAR1A* [2]. In addition, we also detected super-enhancers for cancer-related genes previously described in other cancers such as *TBX3, MAML3* and *ALK* [9, 10, 19] (Figure 1d, Supplementary Table 1).

Next, we performed gene ontology (GO) analysis of all super-enhancer target genes. We identified gene enrichment for GO terms in functional classes such as sterol homeostasis, regulation of transcription, regulation of cell differentiation, cell migration and epithelial to mesenchymal transition (EMT) (Figure 1e, Supplementary Table 3).

### Super-enhancer target genes are predictive for patient survival

Given known oncogenic properties of many genes regulated by super-enhancers in ACC (Figure 1d-e), we addressed the question as to whether expression of such genes could serve as a good clinical biomarker. We performed survival analysis based on gene expression and clinical data from *The Cancer Genome Atlas* (TCGA) [2] (*N* = 76). To build a predictive model, we used the top 500 genes regulated by super-enhancers (highest mean super-enhancer score over all ACC samples). To avoid over-fitting, we used LASSO regularization [20] in a linear model (Methods). By assessing the feature importance (Methods), we identified the *LMNB2* gene coding for Lamin B2 as a stable predictor for patient survival (Figure 1f).

To validate the predictive performance of our model with the top 500 super-enhancer-regulated genes, we binarized the dependent variable to predict groups of poor and good outcome and benchmarked it against the model based on the molecular subgroups C1A/C1B currently used in clinics [4]. In the nested cross-validation setting, our model based on genes regulated by super-enhancers achieved a concordance score of 0.7688 (log rank test *p*-value = 1.46 · 10^−5^) outperforming the molecular subgroup C1A/C1B classification (concordance score 0.738, log rank test *p*-value = 8.57 · 10^−5^) (Figure 1g).

### The super-enhancer landscape reflects known molecular subgroups

Unsupervised study of aggregated H3K27ac scores of 3,742 ACC super-enhancers with principal component analysis (PCA) revealed that the super-enhancer landscape significantly differed between the molecular subgroups C1A and C1B (Figure 2a, two-sided Wilcoxon rank sum test on PC1, *p*-value = 3.1 · 10^−5^). We also tested for associations between principal components and other clinical features and the CIMP phenotype, but no other association was significant after the false discovery rate (FDR) correction (Supplementary Table 2).

**Figure 2:**
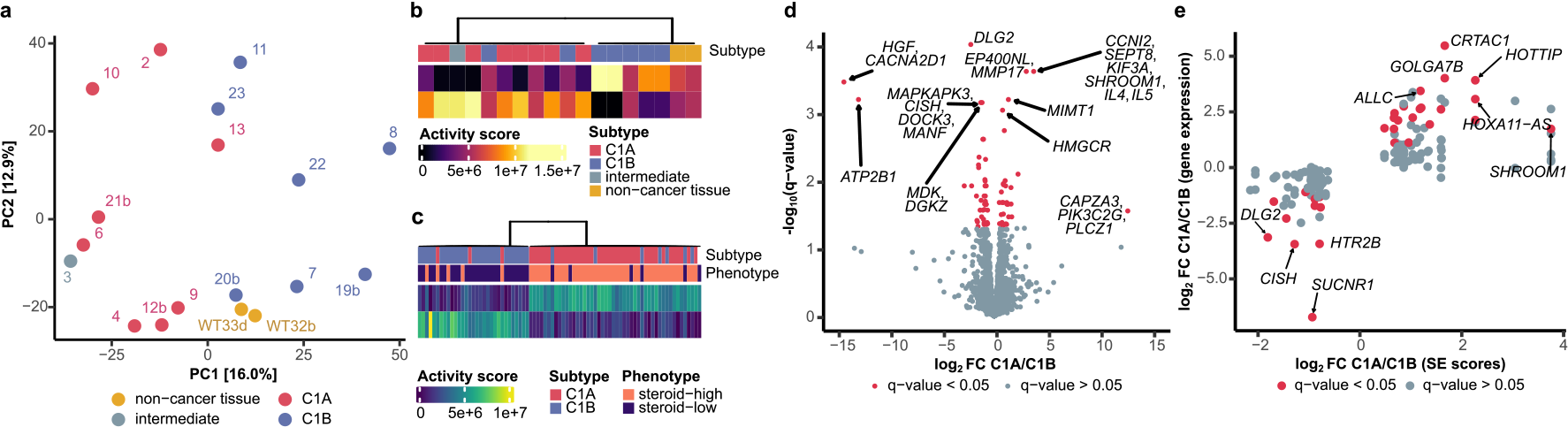
The super-enhancer landscape reflects molecular subgroups of ACC. **a**, PCA of all super-enhancer region scores (*N*=3,742) that were identified in ACC samples (*N*=16) and samples of non-cancer tissue (*N*=2). The samples are labeled according to their sample name, the color reflects the molecular subgroup. **b**, Output of NMF on all super-enhancer scores (*N*=3,742) in all patient samples (*N*=16). The colors indicate the gene expression subgroup. **c**, Output of NMF on gene expression of super-enhancer targets (*N*=2836). Gene expression data are taken from the TCGA ACC dataset (*N*= 76). The colors indicate the molecular subgroups and steroid phenotype. The rows of the matrices in (**b**) and (**c**) correspond to the signatures identified by NMF, the activity score indicates the weight given to the respective signature. **d**, Volcano plot illustrating super-enhancer score differences between the two molecular subgroups. Colored in red are regions with significantly different super-enhancer scores (*q*-value <0.05 from edgeR’s likelihood ratio test [22]). **e**, Scatterplot of super-enhancer target genes with differential activity scores between the subgroups (*N*=161). The x-axis shows the log_2_ fold change C1A/C1B of the super-enhancer score, the y-axis the log_2_ fold change of the gene expression value of the patient cohort. Colored in red are genes with significantly different gene expression values (edgeR’s likelihood ratio test, *q*-value <0.05). Spearman’s correlation 0.78, *p*-value <10^−20^. Genes with FC smaller than -20 or greater than 20 are not shown. FC, fold change.

We further confirmed that the two types of super-enhancer profiles in ACC cells were linked to molecular subgroups C1A and C1B by applying a non-negative matrix factorization (NMF) [21], a signal deconvolution method able to extract hidden signatures. Indeed, the NMF signatures extracted from super-enhancer scores (3,742 regions) of 15 ACC tumors and two non-cancer adrenal grand samples reflected the two molecular subgroups (Fisher’s exact test, *p*-value = 0.007, Figure 2b).

We further examined whether subgroup specific differences were also present on the expression level of super-enhancer target genes. For this, we applied NMF on trascriptomics data from our cohort and in the TCGA ACC dataset (*N* = 76) using as input expression values 2,836 genes associated with ACC super-enhancers. The NMF signatures corresponded to the two molecular subgroups both in the TCGA ACC dataset (Fisher’s exact test, *p*-value = 2.6 · 10^−11^, Figure 2c) and in our cohort (Fisher’s exact test, *p*-value = 0.03, Supplementary Figure S1).

To characterize specific differences in super-enhancer activity between molecular subgroups, we performed differential analysis of super-enhancer scores (Methods). We identified 76 super-enhancer regions in which the super-enhancer scores significantly differed between the C1A/C1B subgroups after FDR correction (Figure 2d). In total, the 76 differential regions were associated with 161 genes, 31 of which we identified as TFs and/or oncogenes. Gene expression analysis indicated that the subgroup specific differences were also present on the gene expression level in both our patient cohort, as well as in the external TCGA ACC dataset (Figure 2e, Supplementary Table 4).

### Subgroup specific core regulatory circuitries (CRCs) in ACC

Transcriptional CRCs are networks of master transcription factors determining cell identity that auto- and cross-regulate each other in a feed-forward manner via binding to superenhancer regions [12, 23]. Based on the analysis of transcription factor binding motifs in detected super-enhancers [9, 24], we predicted CRCs in 15 ACC tumor samples and two non-malignant adrenal tissue samples (Figure 3a).

**Figure 3:**
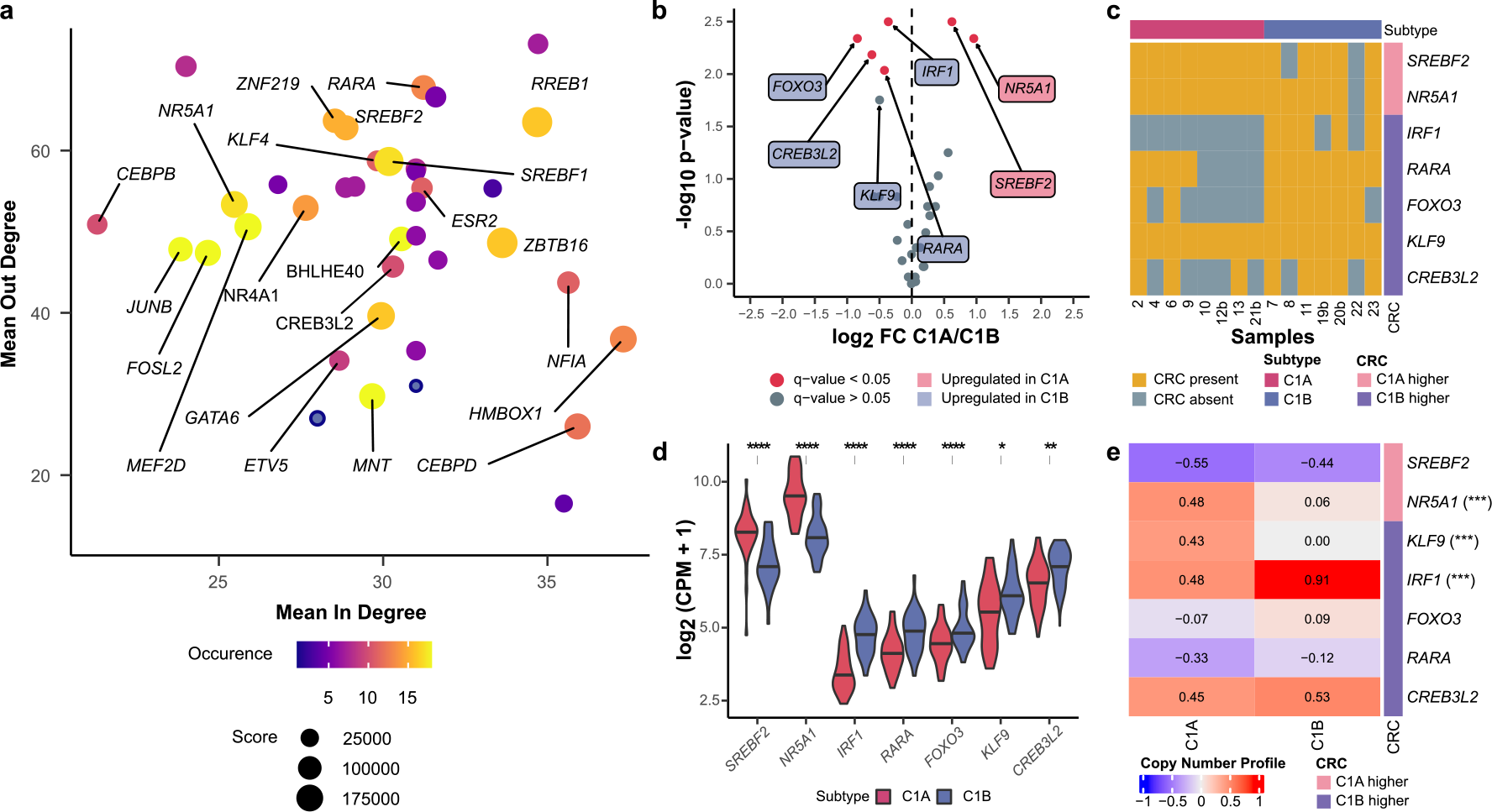
Core regulatory circuitries of ACC. **a**, Scatter plot of CRC associated transcription factors identified by COLTRON in all patient samples (*N*=17). Mean in degree refers to the mean number of transcription factors that bind to a super-enhancer node, mean out degree to the mean number of super-enhancer that each respective transcription factor binds to. The size represents the super-enhancer score and the color the number of samples each CRC was identified in. CRC transcription factors identified in more than half of all samples are labeled with the gene name. **b**, Analysis of CRC transcription factors with differential super-enhancer activity scores between the C1A and C1B molecular subgroups. CRCs that occurred in at least half of the samples of the respective subgroups were tested. Labeled are transcription factors with a two-sided Wilcoxon signed-rank *p*-value <0.05, red dots indicate a *q*-value <0.05. The color in the box signify in which subgroup the super-enhancer scores were higher. **c**, Predicted CRC membership for genes with differentially active super-enhancers identified in (**b**) in patient samples from the two molecular subgroups. **d**, Gene expression of transcription factors identified in (**b**). Gene expression values were taken from the TCGA ACC dataset (*N*= 90). *q*-values were derived from edgeR’s exact test, **** *q*-value <0.0001, ** *q*-value <0.01, * *q*-value <0.05 **e**, Copy number aberrations (CNA) of the transcription factor genes identified in (**b**). Data were taken from the TCGA ACC dataset (*N*= 90) and averaged across patients of the same molecular subgroup. Copy number profile: -1, genomic loss; 0, no change; 1, genomic gain. Indicated with (***) are CRC genes with significantly different copy number profiles between the molecular subgroups, *i*.*e*., *q*-value <0.01 derived from Fisher’s exact test. FC, fold change.

We investigated differences in the activity scores of super-enhancers that regulate CRC transcription factors between the C1A and C1B molecular subgroups. We found that *SREBF2* and *NR5A1* had enriched activation scores in C1A, while *IRF1, FOXO3, RARA, CREB3L2* and *KLF9* were enriched in C1B (two-sided Wilcoxon rank sum test, *p*-value *<* 0.05 and *q*-value *<* 0.1, Figure 3b,c). Gene expression analysis showed that the differences were also present in the expression values of CRC transcription factors, both in our patient cohort and the TCGA ACC dataset (Figure 3d, Supplementary Table 4). Although the majority of CRC transcription factors are shared between C1A and C1B, our findings suggest the existence of several genes that may serve as potential master transcriptional regulators of these molecular subgroups.

Next, we examined gene copy number status of *NR5A1* and other potential drivers of transcriptional programs of C1A and C1B in the two molecular subgroups. We observed statistically significant correlations between molecular subgroups and gene copy numbers of 3 CRC transcription factors: in C1A, a frequent copy number gain of *NR5A1* and, laying 55 Mb apart from it on chromosome 9, *KLF9*; in C1B, a frequent copy number gain of *IRF1* (Figure 3e) (*NR5A1*, Fisher’s exact test *q*-value for gain in C1A = 0.005; *KLF9, q*-value = 0.005; *IRF1*, Fisher’s exact test *q*-value for gain in C1B = 0.003). For *NR5A1* and *IRF1*, gene copy number gains mirrored the observed increase in super-enhancer activity (values normalized for the copy number status with HMCan [25]) and gene expression in C1A and C1B, respectively.

A closer look on copy number profiles around the *KLF9* gene, a possible master transcription factor overexpressed in C1B, revealed that *KLF9* was located in close proximity to a region of frequent loss in C1A patients (Supplementary Figure S4). However, this loss occurred downstream of the *KLF9* gene and did not include any known cis-regulatory regions linked to *KLF9* [26]. The fact that gene expression of *KLF9* was observed to be higher in the C1B subgroup, despite copy number gains in the C1A subgroup, indicated that *KLF9* super-enhancer activity had an dominant effect on gene expression.

### Expression of CRC transcription factors predicts molecular subgroups

We validated the connection between CRCs and the molecular subgroups C1A and C1B by analyzing the predictive performance of CRC gene expression values in a binary classification task on the TCGA ACC dataset. The expression level of the seven CRC transcription factors with differential activity scores between the subgroups were used to build a classification model to predict the molecular subgroups C1A and C1B for each sample.

We observed a high classification performance in a leaveone-out cross-validation scheme using the seven transcription factors with differential super-enhancer scores. A random forest classifier achieved an AUC (area under the curve) of 0.921 (Supplementary Figure S3a). Moreover, only a subset of the variables seemed to be truly important. Looking at the frequency at which the variables were selected, we saw that *NR5A1, KLF9* and *FOXO3* were especially predictive for the molecular subgroup (Supplementary Figure S3b).

We also extended the model to include the expression of the transcription factors that are regulated by the CRC transcriptions factors, the extended regulatory network (Methods). With this bigger model, the performance improved significantly to an AUC of 0.99 (Supplementary Figure S3c-d).

### Response of ACC cell lines to treatment with BRD4 and CDK7 inhibitors previously reported to affect expression of super-enhancer target genes

We investigated the impact of agents previously reported to affect expression of genes driven by super-enhancers on three distinct ACC cell lines, NCI-H295R [27], MUC-1 [28] and TVBF-7 [29]. The cell lines were treated with bromodomain and extraterminal (BET) inhibitor JQ1, cyclin-dependent kinase 7 (CKD7) inhibitor THZ1, which had been shown to target super-enhancer associated genes [30]. We also employed mitotane, an adrenolytic DDT derivative and the standard-of-care drug for ACC patients [31], and forskolin, a steroidogenesis stimulator [29].

To investigate potential transcriptional reprogramming of ACC cells upon specific treatments, we performed RNA-seq analysis of the three cell lines treated with mitotane, forskolin, JQ1, THZ1, and the combination of THZ1 and mitotane. We applied principal component analysis on the expression values of the union of CRC genes and genes with super-enhancer scores differential between the two subgroups to develop the C1A/C1B scoring approach and classify cell lines according to their molecular subgroups (Methods, Figure 4a). Using this classification, TVBF-7 was allocated to the C1A subgroup, which is in line with an extraordinary strong sterodogenic phenotype of this cell line including autonomous cortisol secretion [29]. NCI-295R, previously reported to exhibit a high steroidogenic phenotype as well [29], was also assigned to the C1A subgroup. The classification of NCI-295R as C1A was further supported by an analysis of H3K27ac ChIP-seq data of this cell line (Methods, Supplementary Figure S5). The MUC-1 cell line, derived from a C1B tumor (ACC120, [1, 28]), was classified as C1B, however, closely to the C1A subgroup. Of note, MUC-1 was known to have comparably low steroidogenic activity, which could be increased by treatment with forskolin [29].

**Figure 4:**
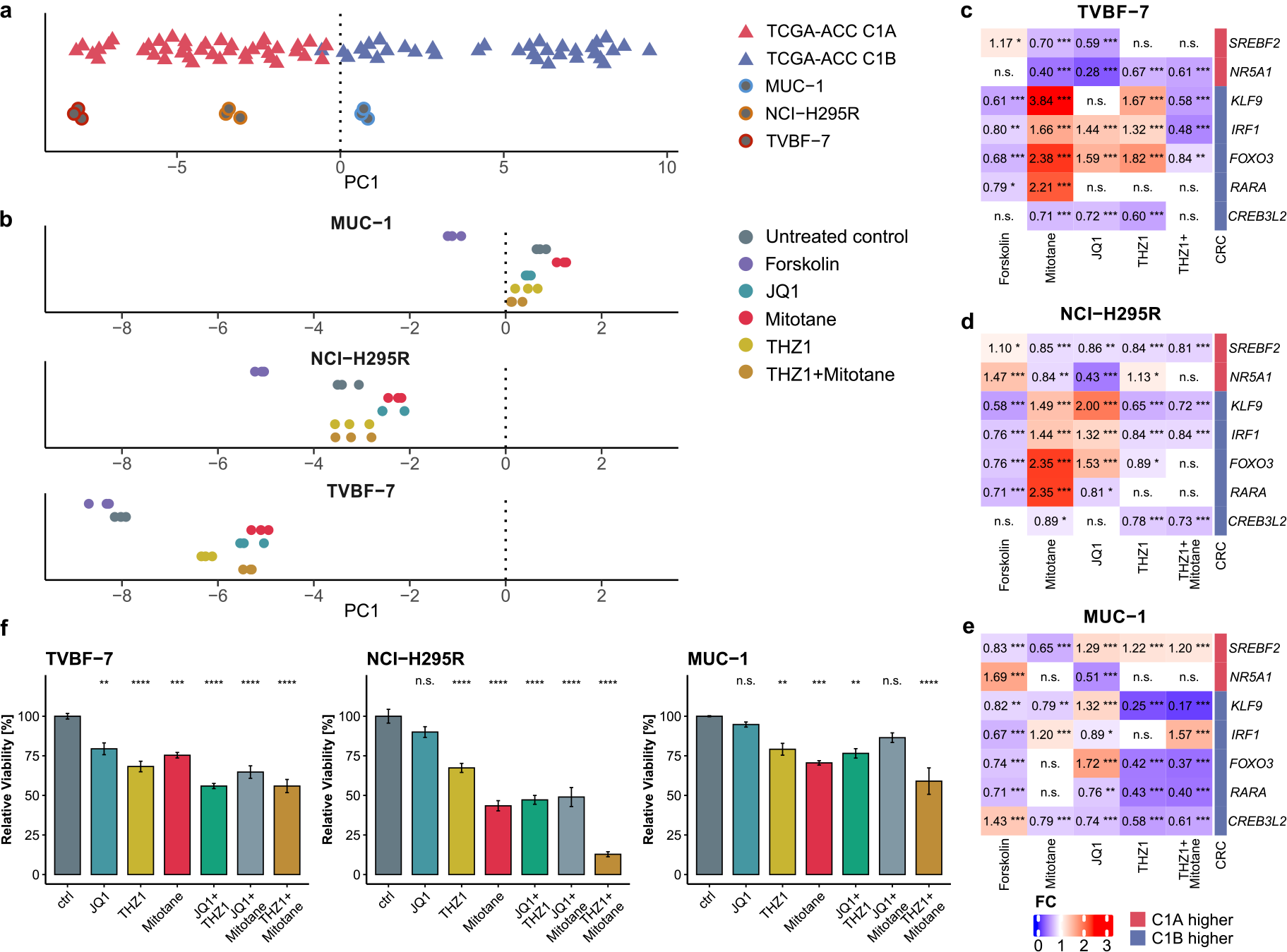
Response of ACC cell lines to treatment with super-enhancer inhibitors. **a**, Projections of the gene expression values on the first principal component (PC1) for TCGA ACC samples (*N*= 74). PC1 explained 18% of variation and discriminated between C1A and C1B molecular subgroups. Gene expression values of genes with differential super-enhancer scores and CRC genes were used. ACC cell line samples (*N* = 9) were projected onto PC1 for subgroup allocation, with the dashed line indicating the border between subgroups C1A and C1B. **b**, Treated and untreated ACC cell line samples (*N* = 54), with color indicating the drug treatment. The dashed line represents the border between subgroups C1A and C1B from (**a**). **c-e**, Differential gene expression of subgroup specific CRC genes after different treatments in cell lines MUC-1 **(c)**, NCI-H295R **(d)** and TVBF-7 **(e)**. The color and values indicate fold change. *P*-values were derived from edgeR’s quasi-likelihood (QL) F-test [22]. **f**, Viablility assays of three ACC cell lines MUC-1, NCI-H295R, and TVBF-7. Barplots represent viability realtive to untreated control (ctrl). *P*-values were derived from Dunnett’s tests. *** *p*-value <0.001, ** *p*-value <0.01, * *p*-value <0.05, n.s., not significant. FC, fold change.

Based on C1A/C1B scores established from gene expression data, we assessed how the C1A/C1B status changed upon treatments that potentially modulated expression of CRC TFs and/or TF-regulated metabolic networks (Figure 4b). As expected, forskolin, an activator of the adenyl cyclase, induced a stronger C1A phenotype in all cell lines. Mitotane, an adrenolytic substance known to inhibit steroidogenesis, showed the opposite effect, inducing a stronger C1B phenotype. JQ1, a BET-inhibitor preventing interactions between the BET proteins and acetylated histones, led to a clear shift from the C1A into the C1B direction in TVBF-7, which had the strongest C1A score. A comparable effect was observed in TVBF-7 after treatment with the CDK7-inhibitor THZ1. Related effects were partly present following treatment with JQ1 in NCIH295R, but not detectable in MUC-1.

We then examined the effects of drug treatment on the expression of the subgroup specific CRC TFs (Figure 4c-e). While forskolin strongly increased expression of *NR5A1* in MUC-1 and NCI-H295R, the response in TVBF-7 was less prominent likely because TVBF-7 already possessed the highest level of basal *NR5A1* expression (Supplementary table 10). Mitotane decreased expression of *NR5A1* in NCI-H295R and TVBF-7, while also increasing the expression of CRC genes that were associated with subgroup C1B, except *CREB3L2*. In TVBF-7, both JQ1 and THZ1 had a similar effect on CRC genes as mitotane. In NCI-H295R, only JQ1 increased expression of most C1B CRC genes. The up and downregulation of subgroup specific CRC genes reflected the cell transcriptional reprogramming along the C1A/C1B axis described above. Interestingly, the combination of THZ1 and mitotane decreased gene expression of almost all CRC genes in all cell lines.

The impact of the treatments on cell viability was to a great extent dependent on the cell line and was decoupled from the drug effects on the transcriptional reprogramming (Figure 4f). Overall, NCI-H295R showed the highest decrease in viability after treatments, followed by TVBF-7, while MUC-1 demonstrated the lowest response. JQ1 alone had a limited impact on cell viability, but when combined with THZ1 or mitotane, a decrease in cell viability was observed in all cell lines. Interestingly, THZ1 applied as single agent demonstrated significant anti-tumoral effects in all cell lines, and the combination of THZ1 and mitotane was particularly effective in NCI-H295R, with an average decrease of around 90 % in cell viability.

### The super-silencer landscape of adrenocortical carcinoma

As we observed that H3K27ac and super-enhancer landscape reflected ACC molecular subgroups, we set up to investigate whether landscapes of other chromatin marks showed differential patterns between C1A and C1B.

Super-silencers, also called H3K27me3-rich regions (MRRs) [32], are large genomic regions characterized by enrichement of the repressive mark histone H3 lysine trimethylation (H3K27me3) deposited by PRC2 (Figure 5a and Methods). To profile super-silencers in ACC, we generated and analyzed H3K27me3 ChIP-seq data for 17 ACC patient samples and 4 samples of non-cancer tissue of the adrenal gland of ACC patients. A total of 1,382 super-silencer regions were detected in at least two samples for which we calculated a super-silencer activity score (Supplementary Table 5).

**Figure 5:**
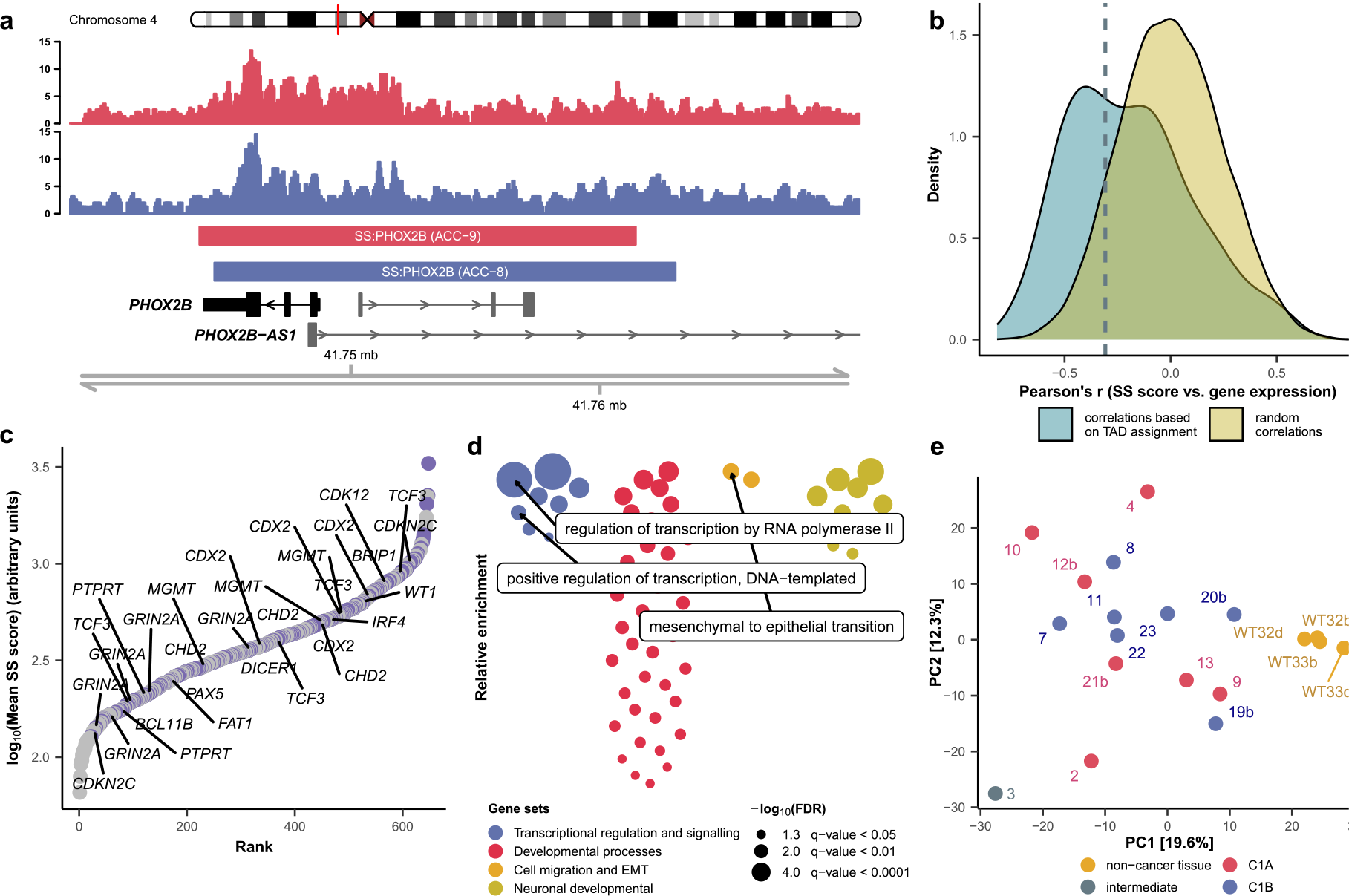
Characterization of the super-silencer landscape of ACC. **a**, An example of a super-silencer associated with the *PHOX2B* gene detected using the H3K27me3 ChIP-seq signal; ACC samples 9 (red) and sample 8 (blue) are shown for illustration. **b**, Distribution of Pearson’s correlation coefficients between super-silencer scores and expression values of putative target genes showing association between activity of super-silencers and gene expression; Correlations with expression of genes in the TAD neighborhood (violet) and random genes (yellow). The red line indicates the correlation threshold determined using the 10% quantile of the distribution for random associations. **c**, Super-silencers ranked by the mean activity score over all ACC samples. Labeled are tumor suppressor genes associated with super-silencers. Blue circles indicate target genes representing transcription factors and/or tumor suppressor genes. **d**, Gene ontology analysis of super-silencer target genes. The colors indicate functional classes. Highlighted in boxes are GO terms relevant for the ACC pathogenesis. Note that these processes are suppressed by the effect of super-silencers. **e**, PCA of the scores of all super-enhancer regions (*N*=1,279) that were identified in the ACC samples (*N*=17) and non-cancer tissue samples (*N*=4). The samples are labeled according to their sample name, the color reflects the molecular subgroup. SS, super-silencer.

**Figure 6:**
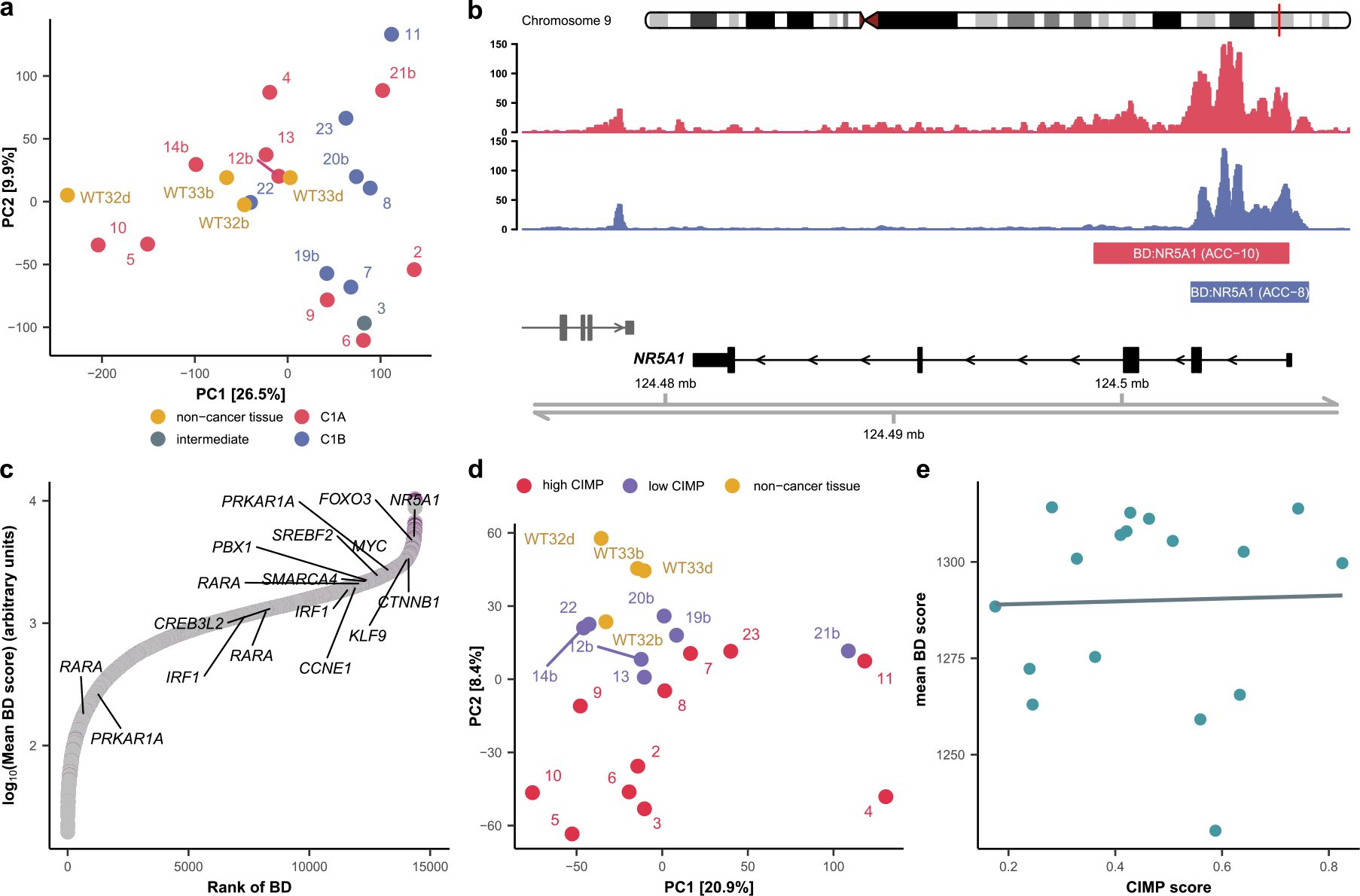
Broad H3K4me3 domains in ACC at active and poised promoters. **a**, PCA of H3K4me3 activity scores of promoter regions (*N* = 39,584) in ACC samples (*N* = 18) and patient samples of non-cancer tissue (*N* = 4). The samples are labeled with their sample names, the color indicates molecular subgroups. **b**, Example of a broad H3K4me3 domain associated with the *NR5A1* gene in ACC. ACC samples 10 (red) and 8 (blue) are shown for illustration. **c**, Broad H3K4me3 domains ranked according to the mean activity score over all ACC samples. Labeled are target genes that were associated with ACC driver genes and important super-enhancer target genes identified in this study. Violet circles indicate target genes of broad H3K4me3 domains for which we also observed super-enhancers. **d**, PCA of the activity scores of broad H3K4me3 regions *N*=14,381 in all patient samples (*N*=22). The color indicate the CIMP phenotype of the ACC samples. **e**, Relationship between the mean activity scores of broad H3K4me3 domains in ACC samples and their respective CIMP score. The blue line indicates the correlation (Pearson’s correlation = 0.03, *p*-value = 0.917). BD, broad H3K4me3 domain.

Using positional information about gene transcription start sites (TSSs) and correlation between super-silencer scores and gene expression, we could assign 648 ACC super-silencer regions to their putative target genes (Figure 5b and Methods). A large number of genes located within the super-silencer regions and regulated by super-silencers represented transcription factors and/or tumor suppressor genes (TSG) (Figure 5c). Among such genes, we observed *MGTM*, encoding a methyltransferase which plays an important role in DNA repair after DNA alkylation, *WT1, PTPRT* and *PHOX2B* all coding for proteins with known tumor-suppressor properties. GO analysis on super-silencer target genes revealed their involvement in the repression of developmental processes (Supplementary Table 7). In particular, high enrichment scores were also observed for regulation of transcription and mesenchymal to epithelial transition (Figure 5d).

Unsupervised analysis of all super-silencer scores using PCA (*N* = 1, 382) showed that the super-silencer landscape differed significantly between the non-cancer tissue and ACC patient samples (two-sided Wilcoxon rank sum test on PC1, *p*-value = 5.2 · 10^−4^, Figure 5e). Associations between principal com-ponents and the molecular subgroups C1A and C1B, clinical variables or the CIMP score were not significant after FDR correction (Supplementary Table 6).

We observed that differences in super-silencer landscape between cancer and non-cancer tissues could be linked to differential expression of *EZH2* coding for a catalyic subunit of Polycomb Repressive Complex 2 (PRC2) mediating deposition of H3K27me3 [33]. It was previously shown that *EZH2* was over-expressed in ACC. Moreover, the excessive *EZH2* expression was related to proliferation and cancer aggressiveness [34]. In our analysis, we observed strong negative correlation between the first principal component of super-silencer scores and *EZH2* expression (Pearson correlation = -0.63, *q*-value = 0.072), driven by the low expression of *EZH2* in the non-cancer tissue samples. The correlation was not significant when only considering ACC samples (Supplementary Table 6). We further evaluated correlation between the principal components of super-silencer scores and gene expression of other subunits of the PRC2 complex (*EZH1, SUZ12, RBBP4, EED*), none of which were significant after FDR correction (Supplementary Table 6).

### Characterization of broad H3K4me3 domains in ACC at active and poised promoters

To study the chromatin state of active and poised promoters, we generated and analyzed ChIP-seq data of trimethylation at the 4th lysine residue of the histone H3 protein (H3K4me3) [35]. The analysis included 18 ACC patient samples and 4 samples of non-cancer tissue of the adrenal gland of ACC patients. We calculated an activity score for each promoter region by summing the ChIP-seq signal around the TSS of each gene. Unsupervised analysis using PCA of all 39,584 promoter scores showed no clear structure in our ACC cohort, no were principal components linked to the ACC molecular subtypes (Figure 5a, Supplementary table 9).

Next, we defined broad H3K4me3 domains (BDs) that span over the TSS into the gene body and act as markers of essential genes [11]. A total of 14,381 BDs were active in at least two samples for which we calculated a BD activity score (Figure 5b, Supplementary Table 8). BDs were assigned to target genes if there was an overlap between the BD and the TSS of a gene. We observed a significant overlap of genes associated with BDs and super-enhancer target genes (Fisher’s exact test, odds ratio = 3.88, *p*-value *<* 10^−20^). Specifically, for 57% of all super-enhancer target genes a BD was also observed (Supplementary Figure S7). Notably, ACC driver genes and most of the CRC transcription factors identified in this study had exceptionally high BD scores (Figure 5c).

PCA of 14,381 activity scores of broad H3K4me3 regions showed no separation between the molecular subgroups C1A and C1B. However, we observed significant correlation between the second principal component and the CIMP score of ACC samples (Pearson correlation = -0.81, *p*-value = 3.57 · 10^−5^). In addition, we also observed significant separation of ACC samples and non-cancer samples on the second principal component (two-sided Wilcoxon rank sum test on PC2, *p*-value = 6.7 · 10^−4^, Figure 5d). The non-cancer samples clustered on the second principal component close to the ACC samples with the lowest CIMP scores.

To explore a possible connection between H3K4me3 and the CIMP phenotype, we studied correlation of the CIMP scores of the ACC samples and averaged H3K4me3 scores of promoter and BD regions. We found no significant correlation between H3K4me3 scores and CIMP in our patient cohort (Figure 5e, Supplementary Figure S8).

## Discussion

The importance of epigenetic modifications in cancer has been elevated in recent years, and epigenetic reprogramming is now regarded as a hallmark of cancer [5]. While the study of epigenetic regulation via DNA methylation has revealed the CIMP phenotype in ACC [6], little was known about other epigenetic regulators, especially histone modifications, in this cancer type. Our work aimed to fill this knowledge gap and focused on the analysis of broad chromatin domains in ACC [11, 32].

First, via the analysis the histone modification H3K27ac, we profiled the super-enhancer landscape in ACC and identified super-enhancer target genes that were associated with ACC pathogenesis. Our results indicated that super-enhancers play a critical role in regulating processes such as steroid phenotype, EMT, and cell migration, which are crucial for ACC pathogenisis. This result was in accordance with what had been described in various other cancers [30, 36] and adds to recent results in ACC cell lines indicating that *β* -catenin binds to *NR5A1* and to maintain differentiation using the adrenal super-ehancer landscape [37]. Moreover, we showed that super-enhancer-regulated genes were important predictors of survival in ACC, with *LMNB2* identified as a potential biomarker [38]. Our study further elucidated that the super-enhancer landscape reflects gene expression subgroups C1A and C1B of ACC and that CRC transcription factors might play a role in defining these subgroups. Treatment of ACC cell lines with drugs previously reported to affect expression of genes regulated by super-enhancers, JQ1 and THZ1, resulted in a heterogeneous response depending on the cell line. Nevertheless, we found evidence of transcriptional reprogramming along the C1A/C1B axis after drug treatment in all cell lines. The expression levels of the super-enhancer-driven genes that we identified as potential master regulators of C1A/C1B molecular subgroups mirrored the transcriptional reprogramming effects upon treatments; this further indicated the relevance of the CRC genes in defining ACC cell identity. We also reported that transcriptional reprogramming effects were decoupled from the effects of the drugs on the cell line viability. These findings however are in accordance with the previously reported specific characteristics of each cell line. NCI-H295R, derived from a primary ACC tumor before signs of metastasis, showed in most cases comparably good therapeutic responsiveness for a wide range of single agents and therapeutic regimens. In contrast, MUC-1 and TVBF-7 represent multi-chemotherapeutic drug resistant cell lines of local and distant metastatic origin [28, 39, 40].

Second, we characterized the super-silencer landscape of ACC by analyzing the histone mark H3K27me3. This analysis showed that there exists no link between the gene expression subgroups of ACC and the super-silencer landscape. However, we saw that the super-silencer landscape differed between ACC cancer samples and samples of non-cancer tissue of the adrenal gland of ACC patients. It is important to note that the noncancer tissue does not necessarily correspond to healthy tissue, as it was also sampled from ACC patients. H3K27me3 is mediated by PRC2, specifically by the catalytic subunit EZH2, which is over-expressed in ACC [34]. The over-expression of EZH2 could be a driving force in defining the super-silencer landscape of ACC. The overall function of super-silencer target genes indicated the importance of long-range epigenetic silencing in the regulation of developmental processes besides the repression of numerous tumor suppressor genes. Specifically, we observed several super-silencer regions associated with the *MGMT* gene (Figure 5c). *MGMT* encodes a methyltransferase which plays an important role in DNA repair after DNA alkylation. Creemers *et al*. have explored the therapeutic potential of DNA-alkylating agents in ACC cell lines. Since *MGMT* can reverse DNA-alkylation, DNA methylation of the *MGMT* pro-moter can indicate treatment effectiveness [42]. *MGMT* promoter methylation was reported to be low in ACC cell lines and adrenal tissues [41]. Our results indicate that the effect of super-silencers on the *MGMT* gene might contribute to efficacy of treatment using DNA-alkylating agents in ACC.

Third, we studied the histone mark H3K4me3 and focused on promoters and broad H3K4me3 domains. We found that target genes of broad H3K4me3 domains and super-enhancer target genes show significant overlap and that ACC driver genes had especially high broad H3K4me3 domain scores. Furthermore, we discovered that the broad H3K4me3 domain landscape partially mirrored the CIMP phenotype. Interestingly, we did not find any associations between broad H3K4me3 scores and the CIMP score of ACC patients on a global genomic level. However, we suggest that this connection should be more thoroughly investigated in further analyses. To the best of our knowledge, no connection between histone modifications and the CIMP phenotype was described in ACC patient samples.

Our study has limitations due to the rarity of ACC, leading to a limited sample size. To overcome this, the TCGA database was used when appropriate. The study only considered bulk ChIP-seq data, limiting conclusions about intratumor heterogeneity. Despite these limitations, our study sheds light on the histone modification landscape of ACC, linking the super-enhancer landscape to molecular subgroups and identifying core transcriptional networks. Drug treatment experiments showed potential for certain super-enhancer-affecting molecules in treating ACC, but differences in treatment outcome were observed between subgroups demanding for further investigation. In sum, our study provides the groundwork for epigenetic characterization of ACC tumors and indicates the potential for novel combination treatment strategies for this aggressive cancer.

## Methods

### Patient samples

Patient samples were obtained between 2008 and 2015 in the Cochin hospital (Paris, France), snap-frozen, and stored as described in Assié *et al*. [1]. An expert endocrine pathologist diagnosed the malignancy of the samples and characterized the samples according to the Weiss criteria [43]. In addition, clinical information including age, sex, hormone secretion, mitotic count, Ki67 staining index, ENSAT tumor stage, disease-free survival, and overall survival was retrieved from the ENSAT database [44].

### ChIP-seq experiments

The iDeal ChIP-seq kit (Diagenode, Serang, Belgium) was used to perform H3K27ac, H3K27me3, and H3K4me3 ChIP-seq analysis according to the manufacturers protocol. Supplementary Information contains a more detailed description of the experimental approach and analysis pipelines. As the quality of ChIP-seq differed between the markers, some samples were excluded from final analyses after quality control.

### Patient sample RNA sequencing

For RNA extraction, tumor samples were first powdered under liquid nitrogen. We then used 10–50 mg of tumor samples and *←* 3 · 10^5^ of cell line cells for RNA extraction with the TRIzol Reagent (Thermo Fisher Scientific, Waltham, MA USA) according to manufacturers protocol. The extracted RNA was treated with DNAse (TURBO DNA-free™ kit, Thermo Fisher Scientific) before RNA-seq was performed. Information about the sequencing procedure and the bioinformatic workflow can be found in Supplementary Information.

### Reduced Representation Bisulfite Sequencing experiments

Genomic DNA was extracted using QIAamp Fast DNA Tissue Kit (Qiagen) according to manufacturer protocol. Reduced Representation Bisulfite Sequencing (RRBS) was performed by Integragen SA (Evry, France) with the Diagenode Premium RRBS kit. Information about the sequencing procedure and the bioinformatics workflow can be found in Supplementary Information.

### Computation of the CIMP score

In order to obtain a quantitative value of the CIMP phenotype for subsequent analyses, RRBS data were used to compute a CIMP score. First, CpG probes located in promoters of genes were extracted (“TSS200” in the Illumina Infinium Human-Methylation annotation file). Probes that were found significantly different between low, intermediate and high CIMP (as defined in Zheng *et al*. [2]) with a delta *β*-value *>* 0.2 between low and intermediate samples and between intermediate and high CIMP were selected as CIMP signature probes. The score was then calculated as the sum the *β*-values of these probes.

### Cell line viability assays

NCI-H295R (8000 cells/well), MUC-1 (6000 cells/well) and TVBF-7 (7000 cells/well) cells were seeded in 96-well plates (TPP, Trasadingen, Switzerland) and incubated at 37°C and 5% CO_2_ overnight. Following media were used for the different cells: DMEM/F12 (Thermo Fisher Scientific, Waltham, MA, USA) supplemented with 2% Ultroser G (Pall Biosepra, CergySaint-Christophe, France), 1% Insulin-Transferrin-Selenium and 100 U/ml Penicillin / 100*μ*g/ml Streptomycin (both Thermo Fisher Scientific) for NCI-H295R and Advanced DMEM/F12 supplemented with 10% FBS and 100 U/ml Penicillin / 100*μ*g/ml Streptomycin (all three Thermo Fisher Scientific) for both MUC-1 and TVBF-7. On the next day, media were replaced with 100*μ*l of the respective media containing forskolin and drugs: forskolin, 1*μ*M THZ1 (Apexbio, Houston, TX, USA), 1*μ*M (+)-JQ1 (Selleck Chemicals, Planegg, Germany) and 40*μ*M mitotane (Sigma-Aldrich, St. Louis, MO, USA). DMSO concentration in all wells (including controls) were adjusted to 0.5% (DMSO, Sigma-Aldrich). Following an incubation time of 24 hours, 10*μ*l of 5mg/ml MTT solution (Apexbio) were added per well. After 2 hours incubation with the MTT solution, cells were lysed in sealed plates upon addition of 100*μ*l of 10% SDS / 10mM HCl at room temperature overnight. The absorbance at 570nm and at 650nm for background was measured with the PowerWave340 plate reader (Biotek, Winooski, VT, USA). Assays were repeated to a total of five times for NCI-H295R cells and to a total of three times for MUC-1 and TVBF-7 cells, respectively. Dunnet’s test was used to evaluate the treated cell lines’ vitality to the untreated control.

### Cell line RNA sequencing

NCI-H295R (1 · 10^6^ cells/well), MUC-1 (5 · 10^5^ cells/well) and TVBF-7 (9 · 10^5^ cells/well) cells were plated in 6-well plates (TPP, Trasadingen, Switzerland). On the next day, media were replaced with 3ml of the respective media containing forskolin (Sigma-Aldrich) and drugs: 25*μ*M forskolin, 1*μ*M THZ1, 1*μ*M (+)-JQ1, and 40*μ*M mitotane. DMSO concentration in all wells (including controls) was adjusted to 0.5%. Cells were incubated for 24h. RNA was extracted with RNeasy Mini kit (Qiagen, Hilden, Germany) and treated with DNAse (TurboDNA-free™ kit, Thermo Fisher Scientific) using manufacturers protocols. RNA Sequencing was performed by the Genomics Facility Basel, department of Biosystems Science and Engineering (D-BSSE), Basel, Switzerland. Information about the bioinformatic workflow can be found in Supplementary Information.

### Statistics and Data analysis

Data analysis was performed using R (version 4.1.2) [45]. *p*-values were adjusted to *q*-values using the FDR correction method by Benjamini & Hochberg [46].

### Detection of histone modification markers

We used HMCan (Histone Modifications in Cancer, version 1.44) to detect genomic regions in which histone modifications (H3K27ac, H3K27me3 and H3K4me3) were enriched [25]. HMCan uses a module from the Control-FREEC method [47] to estimate the copy number and explicitly corrects for copy number variation. In addition, HMCan accounts for GC-content bias resulting from the PCR amplification step and masks out known ‘blacklist’ regions in which signal is unreliable independently of the cell line or experimental setup [48]. We further used ChIP-seq inter-sample normalization technique, ChIPIN, to normalize the ChIP-seq signal [49]. We used linear normalization for H3K27ac and H3K4me3, and quantile normalization for H3K27me3 due to differences in specificity of the antibodies. Super-enhancers, super-silencers and broad H3K4me3 domains were detected using LILY [9] which is based on the Rank Ordering Of Super-Enhancers (ROSE) algorithm [12].

For super-enhancers, LILY stitched enhancers within a region of 12.5 kb together while excluding promoter regions (*±*2.5 kb from the transcription start site (TSS)) to identify clusters of enhancers. Next, the clusters were ranked according to their ChIP signal score and a threshold determined by ROSE distinguished super-enhancer from other enhancers. For super-silencers and broad H3K4me3 domains we used a stitching size of 4kb and included promoter regions. In addition, we calculated an H3K4me3 activity score for each promoter region by summing the normalized ChIP-seq signal in a region from *-*1000 bp to +2000 bp around the TSS gene coordinates taken from the Ensembl database [50].

### Normalization of histone modification scores

To avoid false positives, histone modification scores were calculated for regions that presented strong histone modification signals in at least two samples. The activation scores were calculated by summing the normalized ChIP-seq signal in the genome region where the epigenetic modification was located. This was done in all samples, regardless whether a particular regions was also identified in the specific sample. While ChIPIN normalized the overall density profile for the samples around the TSS, there still existed a heterogeneity in the ChIP-seq signal between different regions of the genome. To correct for the differences in the overall signal, we defined a measure called mean noise per base (*mnpb*). It was calculated for each sample individually by first summing the signal of all regions in the genome that did not correspond to a histone modification, enhancer or promoter region identified by LILY. We further excluded blacklist regions that were previously ignored by HMCan. This sum was then divided by the total length of all regions to estimate the *mnpb*. We then subtracted from each activity score the *mnpb* times the length of the respective region. In addition, we used edgeR [22] to normalize the histone activity scores between the different samples using the trimmed mean of M-values (TMM) normalization.

### Identification of target genes

We first assigned super-enhancers and super silencers to topologically-associated domains (TADs). We used TAD information of eight human cell lines that were described by Rao *et al*. [51]. Genes whose TSS fell into the same TAD region as a super-enhancer/super-silencer were associated to the respective super-enhancer/super-silencer; several assignments were possible for each super-enhancer/super-silencer. Second, we calculated the correlation between the histone mark enrichment score of a gene and its corresponding gene expression value using a one sided Pearson correlation test. To determine a correlation value threshold we performed a series of repeated correlation tests of the activation scores with random genes. The threshold for further analysis was defined as the correlation coefficient at the 90% quantile of all values for super-enhancers (threshold = 0.34, Figure 1c) and the 10% quantile of all values for super-silencers (threshold = *-*0.31, Figure 3b). Broad H3K4me3 domains (BDs) were assigned to genes whenever there was an overlap with a TSS. The TSS gene coordinates were taken from the Ensembl database [50]. More than one BD region could be assigned to one gene and more than one gene to each BD region. Transcription factors (TFs) were identified using the list of human TFs by Lambert *et al*. [52]. For oncogenes, the list of COSMIC (including tier 1 and tier 2 data) was used [53].

### Identification of core regulatory circuitries

We used COLTRON [24] as described by Boeva *et al*. [9] to identify core regulatory circuitries (CRCs) in 15 ACC tumor samples and 2 samples of non-cancer tissue. For each candidate gene in a CRC, COLTRON calculated an in-degree that referred to the number of transcription factors that bind to a super-enhancer and an out-degree which represented the number of super-enhancer that each respective transcription factor binds to. COLTRON identified a total of 267 super-enhancer-associated transcription factors in all samples. 146 of these super-enhancer targets were also identified using our super-enhancer identification algorithm, while 121 targets were not identified (80 of which were detected but did not pass the gene expression correlation threshold). As not all CRC genes were identified in every sample we used super-enhancer scores calculated in our analysis to assess super-enhancer score differences between subtypes. We selected CRCs that were identified in at least half of the samples of each respective subgroup and that were detected in our analysis (*N* = 30, Supplementary Figure S2). We then tested differential super-enhancer score using a two-sided Wilcoxon signed-rank test. For further analysis we selected CRC genes with *p*-value *<* 0.05 and *q*-value *<* 0.1. Differences between ACC and non-cancer tissue samples were not significant after FDR correction.

### Differential feature analysis

We used edgeR [22] to test super-enhancer score differences and detect differentially expressed genes. Differential gene expression analysis was conducted on both our patient cohort (*N*=17) and the TCGA ACC dataset (*N*=76) and all cell line samples (*N*=54). Differences in the number of genes tested (Supplementary table 4) arise from different reference gene annotations and filtering by edgeR.

### Unsupervised subgroup assignment

Principal component analysis (PCA) was performed on all histone modification score values after scaling and centering the data. We investigated whether cancer or non-cancer tissue, the molecular subgroups, the CIMP score and other clinical variables contributed significantly to the explained variance of the first two principal components. We performed a two-sided Wilcoxon rank sum test for group assignments and a two-sided Pearson correlation test for numerical values. The tests were adjusted using FDR correction.

### Gene ontology analysis

Gene ontology enrichment analysis for super-enhancer target gene was done using publicly available code and methods described by Gartlgruber *et al*. [10] adapted to the 2021 release of Gene Ontology biological processes. Gene ontology terms with *q*-values < 0.05 were selected.

### Non-negative matrix factorization

Non-negative matrix factorization (NMF) [21] was calculated using the R package Bratwurst [54] and an adapted script based on publicly available code from Gartlgruber *et al*. [10]. NMF decomposes a matrix of observations **V** into an exposure matrix **H** and a signature matrix **W**. This was done for different ranks *k* and 40 random initializations. The ideal factorization rank was determined using metrics including the Frobenius error, mean Amari distance and cophenetic correlation coefficient. Based on this comparison rank *k*=2 was the optimal factorization (Supplementary Figure S1). Since NMF is invariant to column scaling on **W** or row scaling on **H**, we normalized **W** such that each column summed up to 1 and then scaled **H** correspondingly. The **H** matrices were used to compare the *k* signatures with the molecular subgroups.

### Extended regulatory network of CRCs

We used the output from COLTRON which provides both the core transcription factors of the CRCs and the adjacency matrix which contained information about the downstream regulatory network. Based on this, we created a list of targets of core transcription factors for each sample and summarized it in a table that contained the counts of how often each transcription factor was targeted, considering all samples. Finally, we built the extended regulatory network by considering those transcription factors that were targeted in at least one sample.

### Regularized survival analysis

The Lasso Cox model was used to work with censored survival data of the form *y*_*i*_ = (time_*i*_, event_*i*_), where time_*i*_ is the survival time and event_*i*_ indicates if an event was observed for patient *i*(= 1, …, *N*).

In the standard Cox model [55], given the vectors of covariates *X*_*i*_ = (*X*_*i*1_, …, *X*_*ip*_) and the assumption that the hazards are multiplicatively affected by the covariates, the hazard for person *i* at time *t* with baseline hazard *h*_0_(*t*) is given as

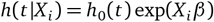

where *β*= (*β*_1_, …, *β*_*p*_) is the vector of regression coefficients.

When considering the likelihood of the event to be observed for subject *i* at time *T*_*i*_, we see that the baseline hazard function *h*_0_(*t*) cancels out and thus does not have to be modeled directly:

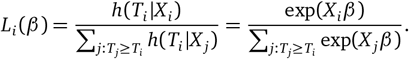

To obtain sparse estimates, the LASSO was combined with the Cox proportional hazards model through a regularized minimization of the log partial likelihood *l*(*β*) = *log L*(*β*) to obtain

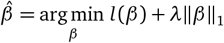

with *λ* being a hyperparameter [20].

### Stability analysis for survival predictions

An important consideration for our use case was the stability of the LASSO-selected non-zero coefficients as stable features were essential for a potential biological interpretation of our results. To test for stability and to estimate the predictive performance, a nested cross-validation scheme was performed. This included an outer Leave-One-Out loop and an inner loop in which 50% of the Leave-One-Out data was randomly sampled. In the inner loop, the optimal hyperparameter *λ* for the regularized Cox proportional hazards model is estimated by cross-validation based on *n-*1 samples and the result is used to identify the non-zero variables. The resulting data matrix is of size *g* × (*s* − 1) with *g* being the number of genes, and *s* the number of samples. This matrix contains the frequency of selection per gene per outer loop in the columns. This procedure was then performed for every constellation of *n -* 1 samples and allowed us to get a better understanding of the stability and predictiveness of the features. The procedure is similar to Stability Selection [56]. The main idea behind it is to repeatedly sub-sample the data and then apply some (arbitrary) variable selection method on it. From this, theoretical properties such as finite sample control for error rates of false discoveries can be derived to guide the process of finding the right amount of regularization [56].

Our procedure differed in the way that we used the outer Leave-One-Out loop to get predictions for the left out observation based on the optimal amount of regularization determined in the inner loop. This was done based on the identification of the highly predictive and stable genes in the inner loop, which were then used to train a model that predicted a hazard for the left out observation. This way we obtained a more realistic estimate of the model performance and additionally got insight into which genes were the most relevant and stable overall.

### C1A/C1B classification of cell lines using PCA

Cell line gene expression data were aligned to TCGA ACC data using joint quantile normalization of FPKM-UQ values. We then performed PCA on the TCGA gene expression data using genes with differentially active super-enhancer scores and subgroup specific CRC TFs; the gene filtering allowed us to discard transcriptional variability due to the potential variation in the amount and composition of tumor microenvironment in the ACC tumors. PCA on the gene expression data from the TCGA ACC database showed that the first principal component (PC1), explaining 18.1 % of the variability in gene expression in the TCGA dataset, was a perfect indicator for the C1A/C1B subgroup allocation. To ensure the validity of this approach for the C1A/C1B group allocation, we projected the gene expression vectors of the in-house patient samples onto the TCGA principal component subspace (Supplementary figure S6); C1A/C1B. status of the in house samples was obtained using an alternative approach based on the expression of marker genes. Except for sample 20b, PC1 perfectly separated C1A and C1B groups validating our molecular subgroup allocation approach. Finally, the projection of the gene expression vectors of the cell lines onto the first principal component (PC) indicated the subgroup allocation of each cell line.

### Super-enhancer landscape of the NCI-H295R cell line

We analyzed H3K27ac ChIP-seq data of the ACC cell line NCI-295R and calculated super-enhancer scores for super-enhancer regions defined in samples of our ACC cohort. After projecting the activity scores on the PCA of the patient samples, we concluded that the super-enhancer landscape of the H295r cell line reflects the C1A subgroup (Supplementary Figure S5a). Gene expression analysis of CRC transcription factors gave further evidence of this allocation (Supplementary Figure S5b).

## Data and code availability

The code used in this study is available at https://github.com/BoevaLab/ACC_chromatin_modifications.

Gene expression (RNA-seq), histone modification (CHIP-seq), and DNA methylation (RRBS) data for the in house ACC tumors can be accessed via EGA`(in_process_for_EGA)`. Cell line data can be accessed via EGA`(in_process_for_EGA)`.

## Acknowledgments

We thank the Genom’IC platform of the Cochin Institute for the sequencing of RNA-Seq experiments.

This work was funded through institutional support from Centre National de la Recherche Scientifique, Institut National de la Santé et de la Recherche Médicale, ATIP-Avenir and the ARC Foundation (ARC-RAC16002KSA-R15093KS), the SIRIC CARPEM and the “Who Am I?” Laboratory of Excellence ANR-11-LABX-0071, funded by the French Government through its Investissement d’Avenir program, operated by the French National Research Agency (ANR-11-IDEX-0005-02) to VB. The project also received funding from the Uniscientia Foundation (keyword tumor model), the Deutsche Forschungsgemeinschaft (project HA8297/1-1), and the CRC/Transregio 205/1 (The Adrenal: Central Relay in Health and Disease) to CH.

## Author contributions

GA, GK and VB designed the study. CH provided the MUC-1 cell line, designed and supervised drug sensitivity experiments. SS and AB provided and characterized the TVBF-7 cell line. GK, MAC, BR and FA carried out the first round of experiments. JH performed sequencing experiments for the tumor samples (ChIP-seq and RNA-seq). IS performed cell line experiments. SG performed data analysis with the help of JM. SG, VB and CH wrote the manuscript. GA, CH, and JB contributed to formulating the hypothesis and editing the manuscript. All authors read and approved the final manuscript.

## Ethics approval

Signed informed consent was obtained from all patients and the study was approved by the local institutional review board (Comité de protection des personnes Ile de France 1, application #13311).

## Extended Data

**Figure S1:**
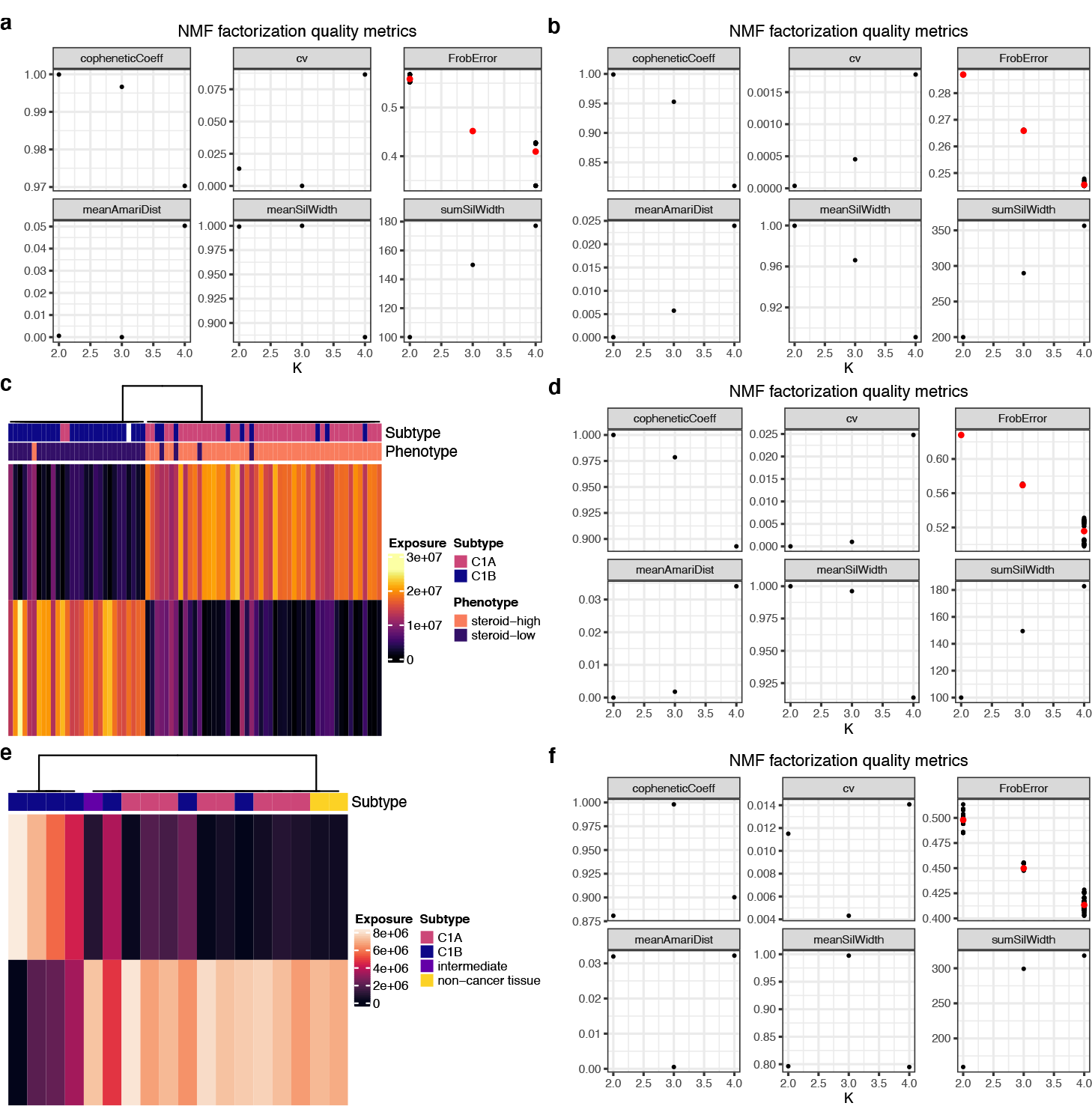
Different metrics to determine the optimal NMF factorization rank for **a**, gene expression data of super-enhancer target genes in the TCGA ACC database (Figure 2b), **b**, all super-enhancer scores (Figure 2c). **c**, NMF of the unthresholded gene expression data of the TCGA ACC database (all genes) and **d**, the respective factorization metrics. **e**, NMF of the super-enhancer target gene expression data of the ACC patient cohort and **f**, the respective factorization metrics. Related to Figure 2.

**Figure S2:**
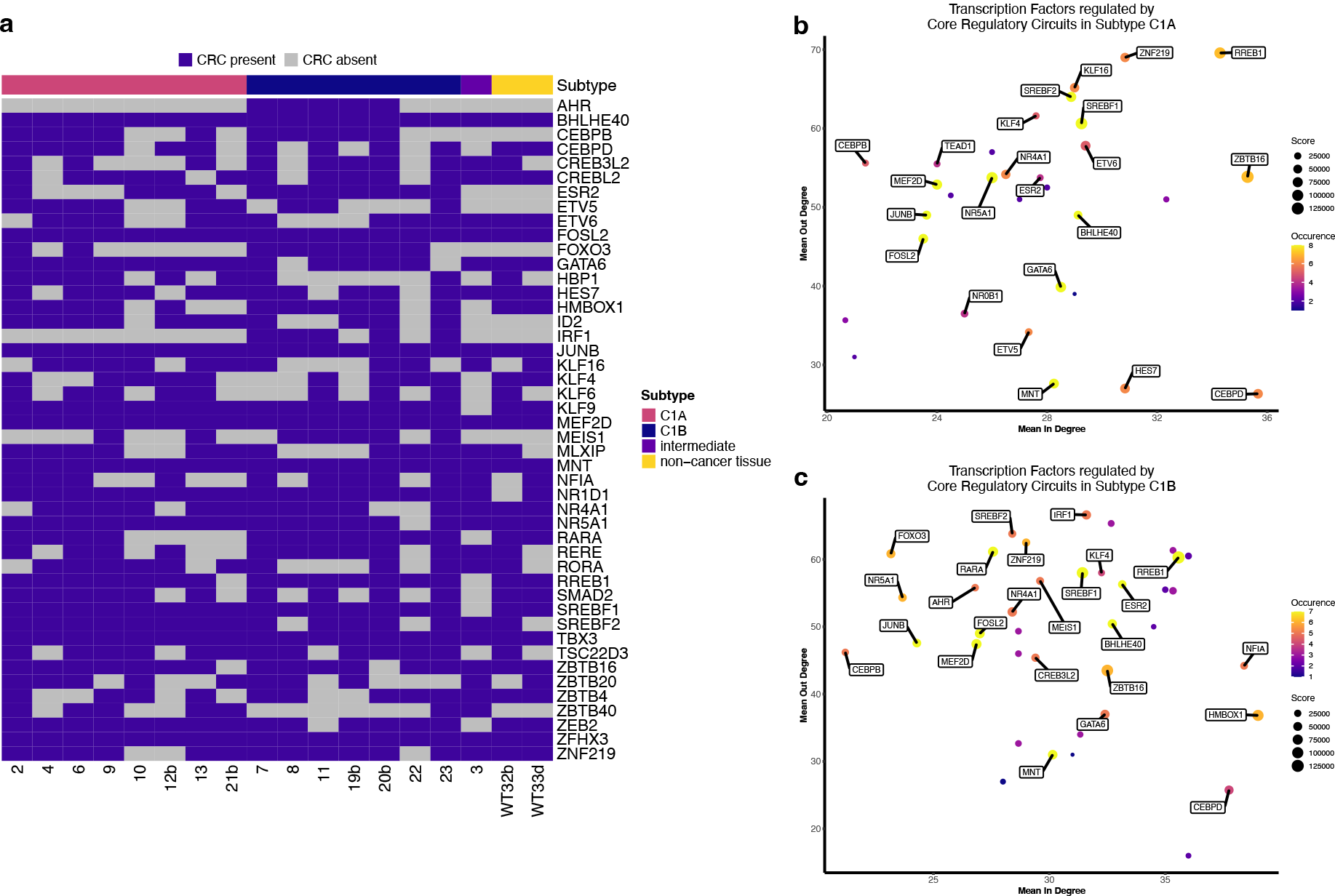
**a**, Predicted transcription factors that participate in CRC in ACC samples (*N*=15) and samples of non-cancer tissue of the adrenal gland (*N*=2). The color indicate the respecitve molecular subgroup. **b-c**, Scatter plots of CRC-associated transcription factors identified by COLTRON in C1A (**b**) and C1B samples (**c**). Mean in-degree refers to the mean number of transcription factors that bind to super-enhancers of a given gene, mean out-degree to the mean number of super-enhancers that each respective transcription factor binds to. The size represents the super-enhancer score and the color the number of samples each CRC was identified in. CRC transcription factors identified in more than half of the samples in each subgroup are labeled with the gene name. Related to Figure 3.

**Figure S3:**
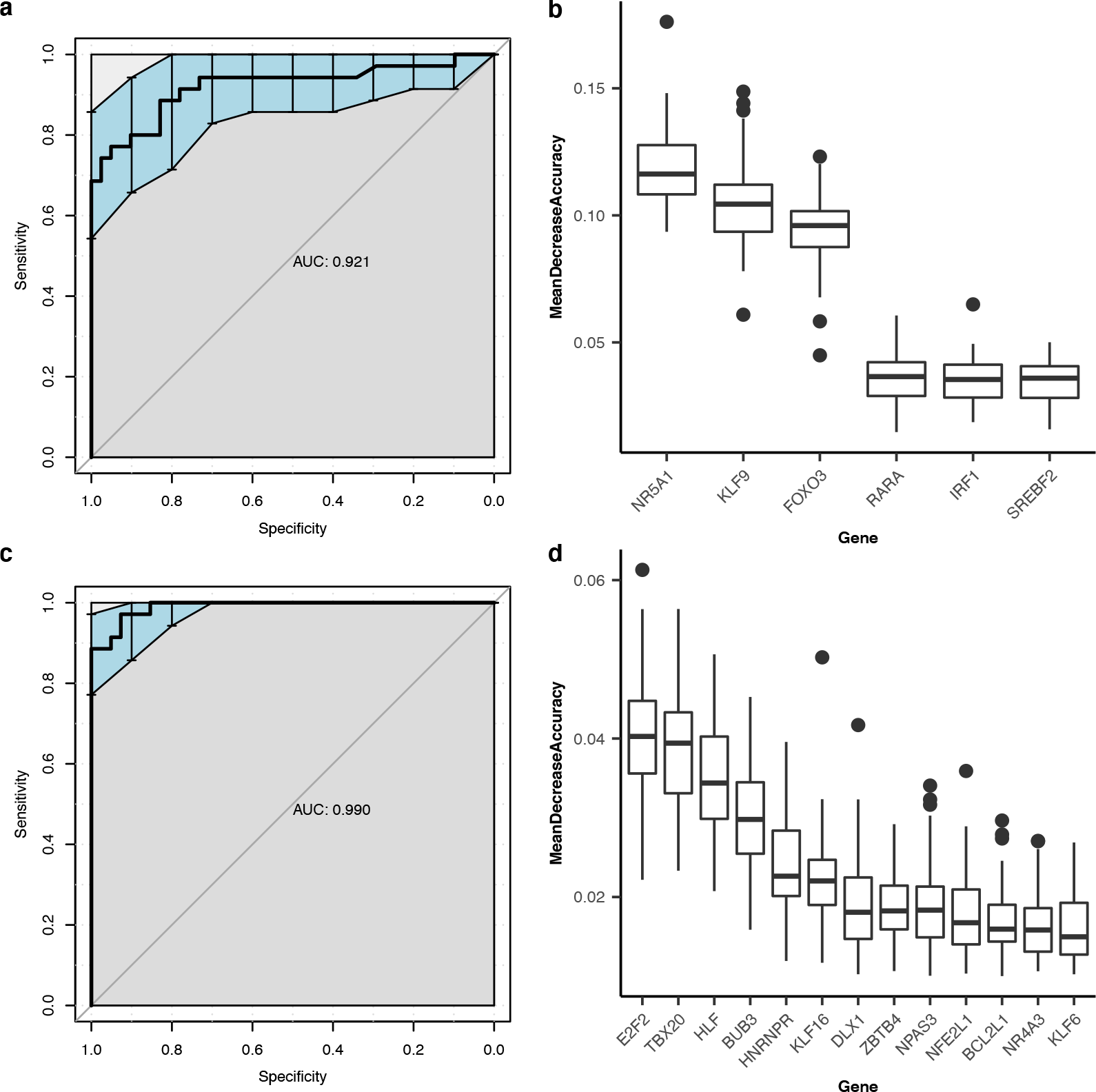
Model performance and variable importance for predicting the molecular subgroup C1A/C1B from CRC data. **a**, The area under the ROC curve (AUC) for the classification task and **b**, the variable importance of the six most important covariates in the model build with the transcription factors that lie at the heart of the respective CRC. (**c**) and (**d**), the AUC and importance for the bigger model that uses the extended regulatory network. Related to Figure 3.

**Figure S4:**
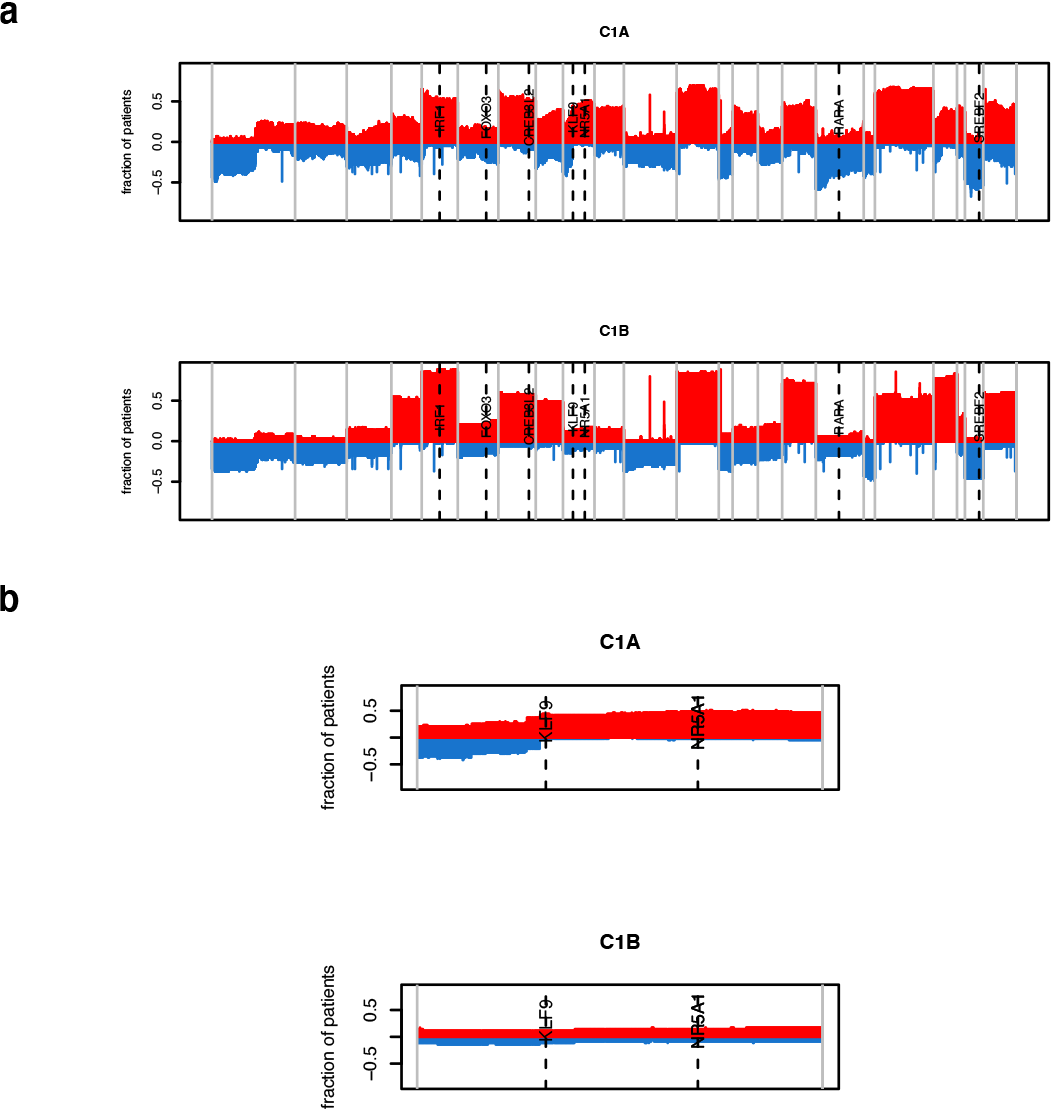
Pattern of copy number alterations (CNAs) in C1A and C1B ACC samples from the TCGA ACC database (*N*=90). **a**, CNA profiles in all chromosomes (separated with vertical grey lines). Indicated with dashed lines are genomic locations of CRC transcription factors identified in this study. **b**, CNAs in chromosome 9 in C1A and C1B samples; indicated with dashed lines are the *KLF9* and *NR5A1* genes. Related to Figure 3e.

**Figure S5:**
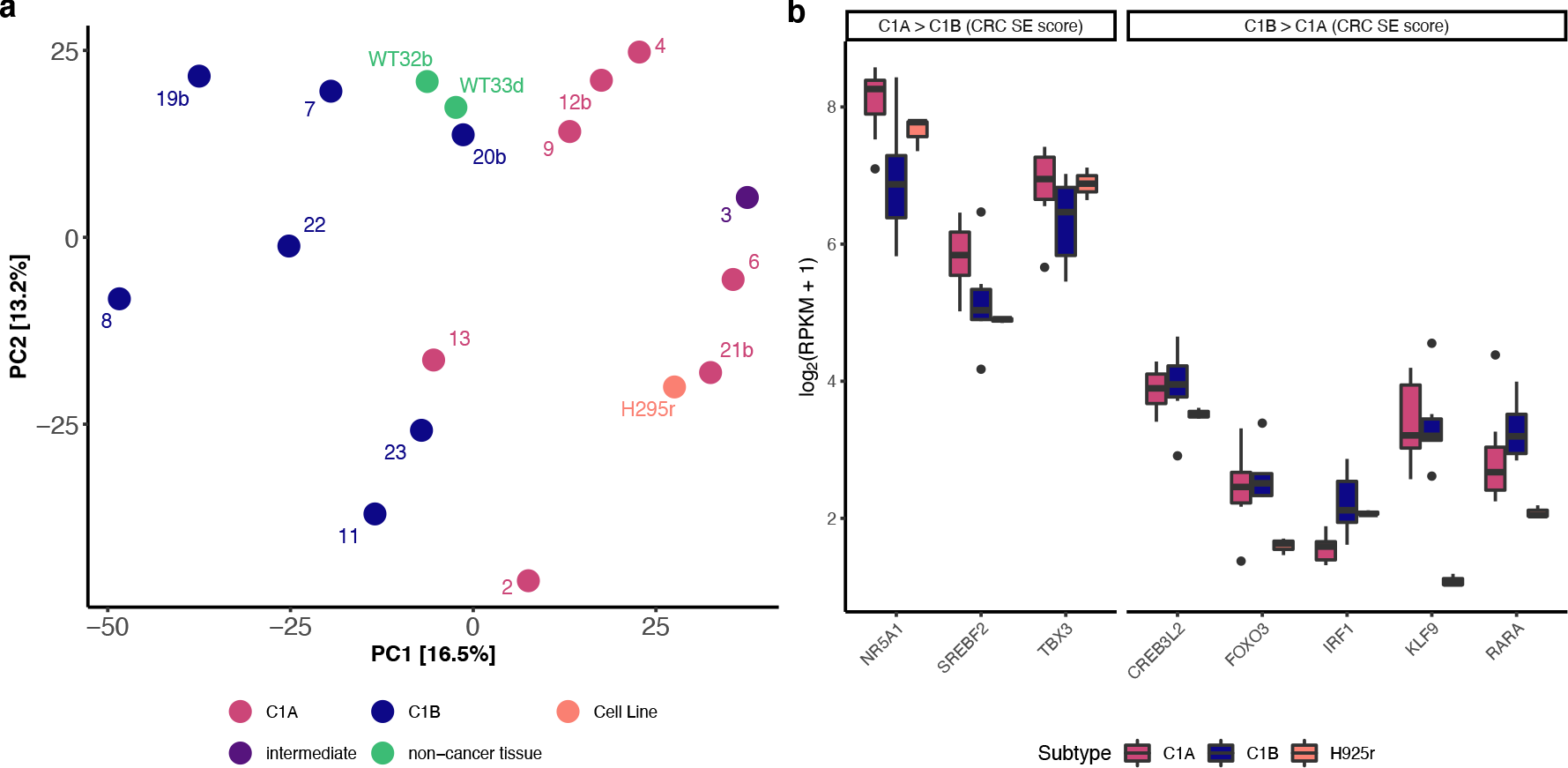
Super-enhancer landscape of the H295r cell line. **a**, Projection of super-enhancer scores calculated for the H295r cell line (*N*=1) on the PCA of super-enhancer scores of the patient samples (*N*=17). **b**, Gene expression analysis of CRC transcription factors identified in this study of ACC patient samples (*N*=15) and the H295r cell line (*N*=3). The RPKM values expression values were jointly normalized using quantile normalization. Related to Methods.

**Figure S6:**
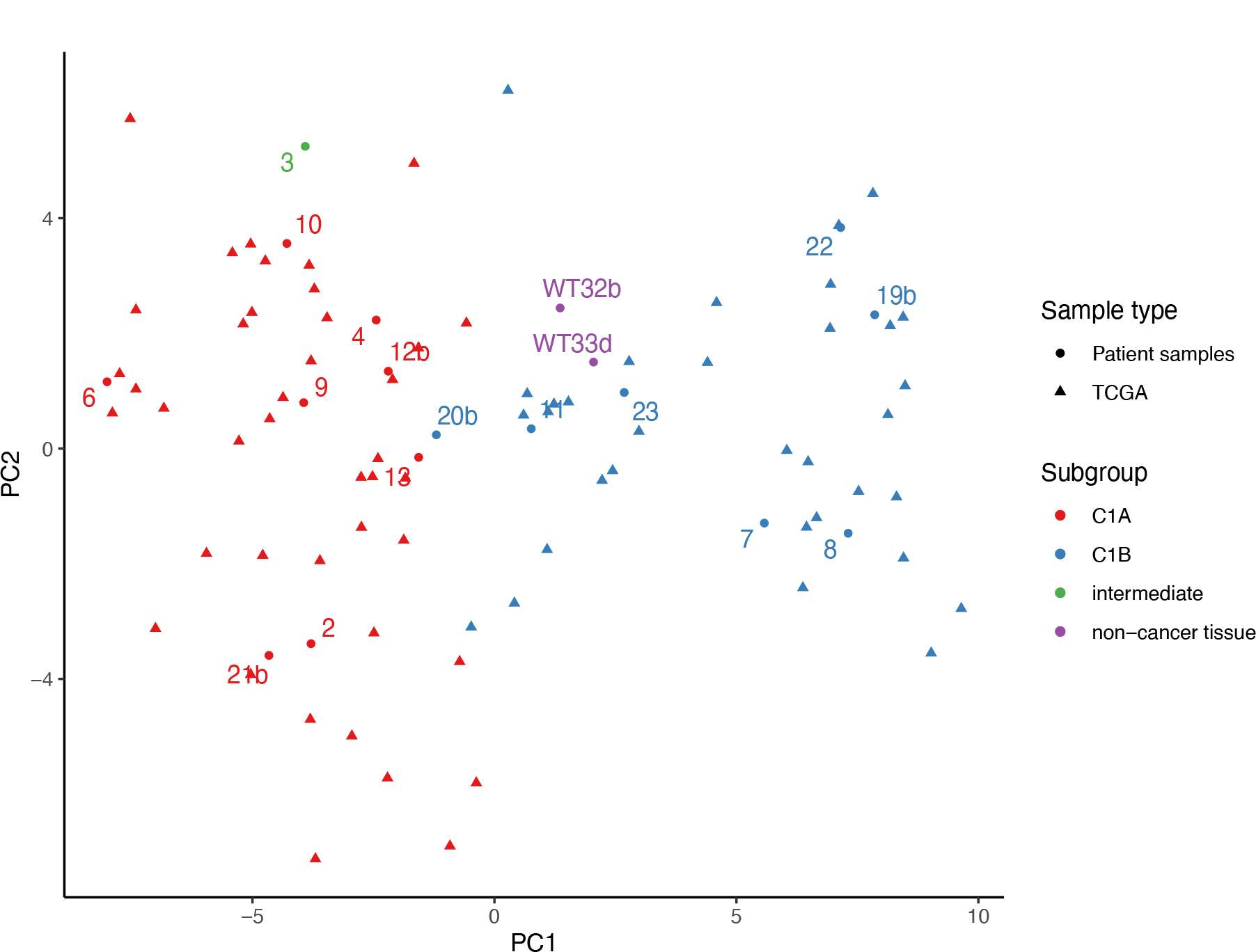
PCA of expression of ACC patient and TCGA samples. Plot of the first two principal components of gene expression of TCGA ACC samples (*N*= 74). Gene expression of genes with differential super-enhancer scores and CRC genes was used. ACC patient samples (*N*= 18) were projected onto the first two principal components for the molecular subgroup confirmation. Related to Figure 4.

**Figure S7:**
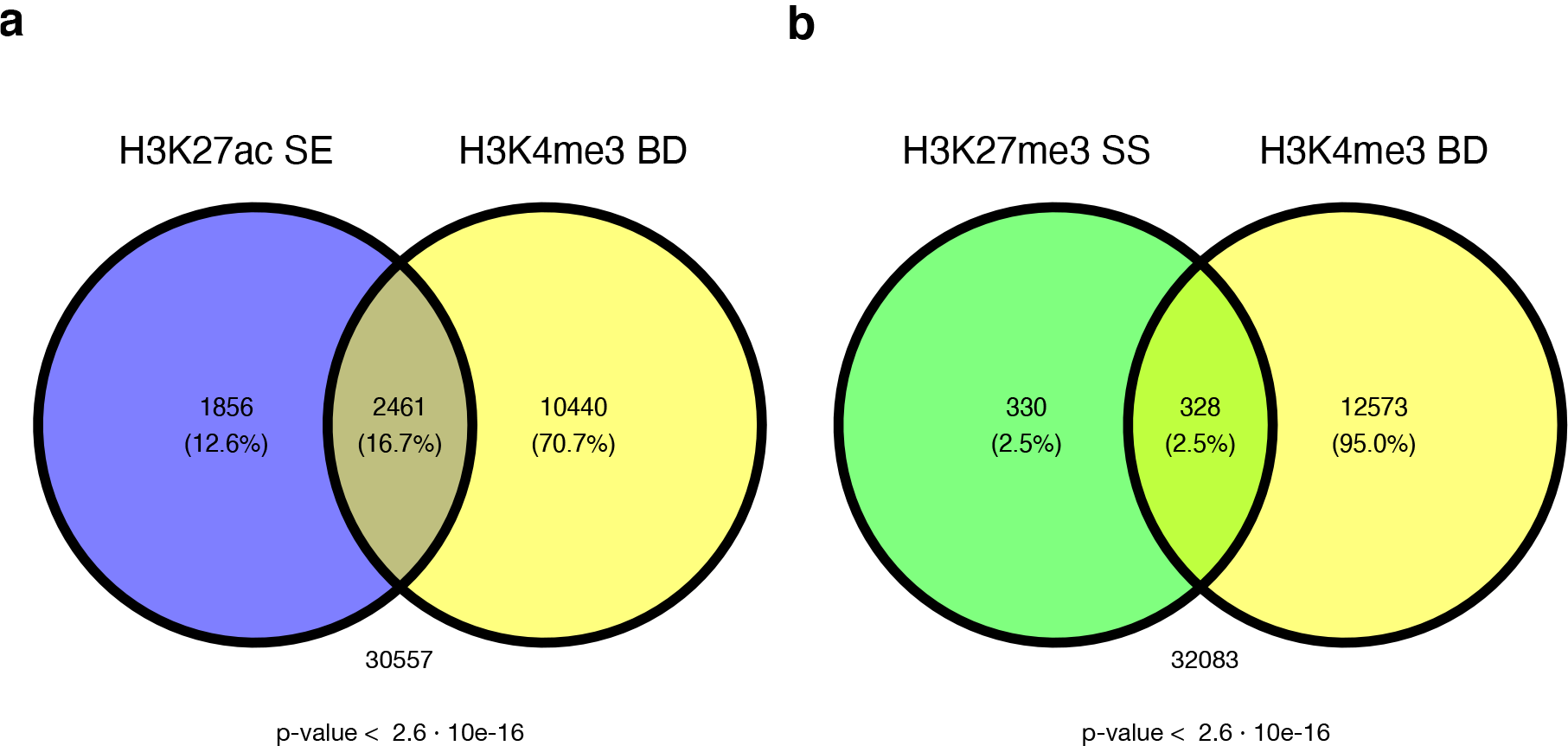
Overlap of gene targets of **a**, H3K27ac super-enhancers and broad H3K4me3 domains (Fisher’s exact test; odds ratio = 3.88, *p*-value <10^−20^) and **b**, H3K27me3 super-silencers and broad H3K4me3 domains (Fisher’s exact test: odds ratio = 2.54, *p*-value <10^−20^). Note that the overlap is based on observed gene targets which does not directly imply an overlap of genomic regions of the histone modifications. Fisher’s exact tests were calculated considering the set of all protein coding and pseudogenes (*N*=45314) taken from [50]. Related to Figure 6.

**Figure S8:**
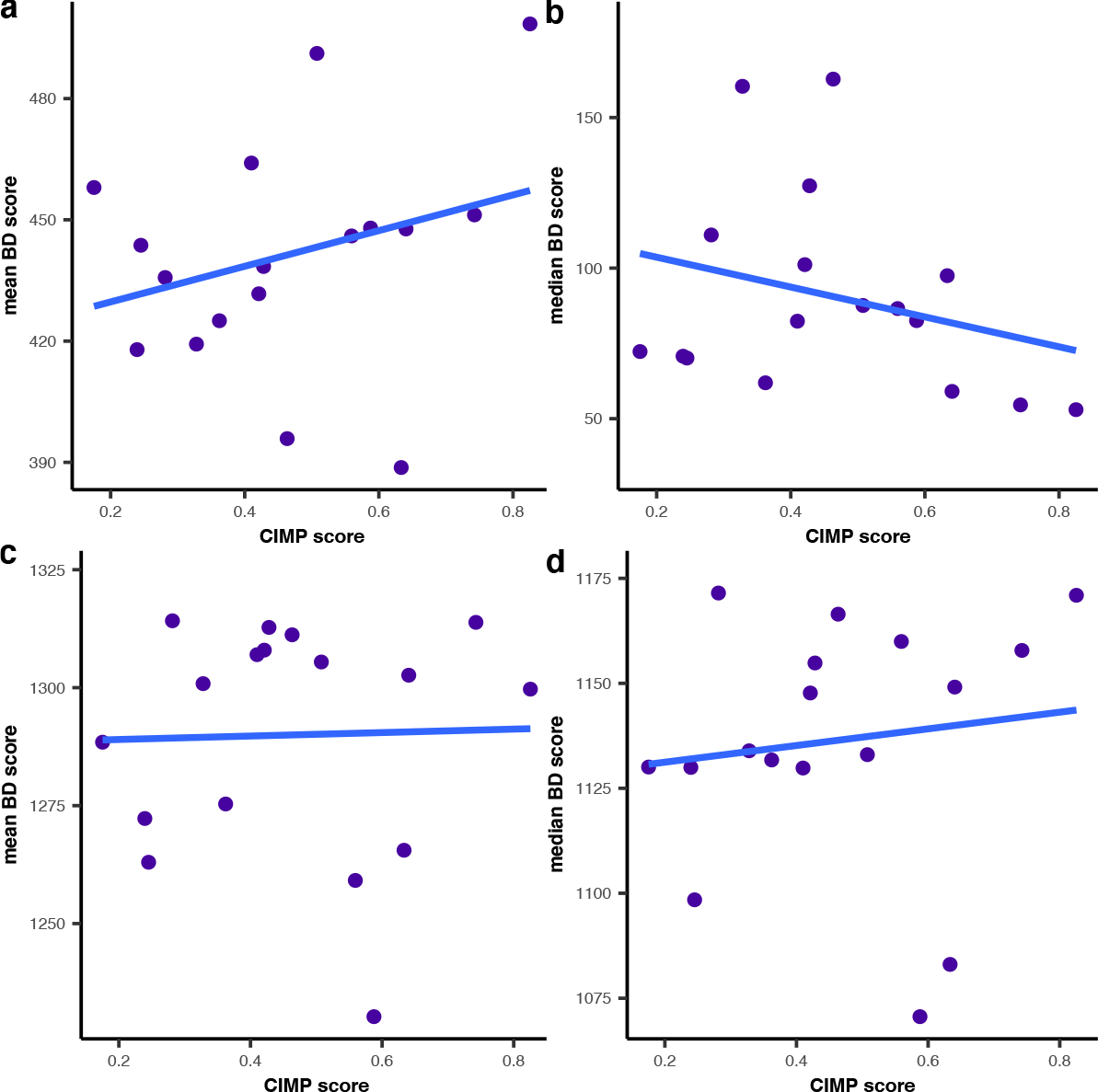
Relationship between H3K4me3 scores and CIMP score of ACC patient samples (*N*=17). **a** and **b**, the relationship between the tumor sample CIMP scores and the mean (**a**) or median (**b**) H3K4me3 scores around gene TSSs; H3K4me3 scores are based on ChIP-seq signal around TSSs of all genes. **c** and **d**, the relationship between the tumor sample CIMP scores and the mean (**c**) or median (**d**) H3K4me3 scores in the broad H3K4me3 domains. The blue line indicate the regression line between the two scores, all correlations were not significant (*p*-value *>* 0.05). Related to Figure 6.

## Supplementary Information

### ChIP-Seq experiments

H3K27ac, H3K27me3 and H3K4me3 chromatin immunoprecipitation (ChIP) were performed using the iDeal ChIP-seq kit for histones (Diagenode) following to the manufacturers protocol. The antibodies used were: ab4729 (rabbit polyclonal, Abcam) for H3K27ac, C15410195 (rabbit polyclonal, Diagenode) for H3K27me3 and C15410003 (rabbit polyclonal, Diagenode) for H3K4me3. Shortly, 20 to 35 mg of tissue were first grinded using Dounce homogenizer and cross-linking was performed using 1% formaldehyde for 10 min. The reaction was ended through addition of glycine (125mM final) for 5 min at room temperature. Chromatin was sheared after addition of lysis buffer using a BioruptorPico (Diagenode) in order to obtain fragments of *←* 300 bp in average. 2% of sheared chromatin was used as input control. IPs were conducted overnight at 4 °C on a rotating wheel in presence of 1*μg* of H3K4me3 antibody or 2*μg* of H3K27ac antibody or H3K27me3 antibody. After 4 washing and elution of the chromatin, the cross-linking was reversed by incubating chromatin 4 h at 65 °C with proteinase K. Finally, DNA was isolated using Ipure kit (Diagenode). ChIP efficiency was validated by qPCR using SensiFAST™ SYBR No-ROX Kit (Bioline) on specific genomic regions depending on each antibody’s targets. The list of primers is available upon request. Reads were aligned to the human reference genome hg38 using BWA v0.7.17-r1188. Low mapping quality reads (Q <20) were filtered out and duplicate reads were removed using samtools v1.758.

### RNA extraction and sequencing of patient samples

Total RNAs extractions were performed using TRIzol Reagent (Thermo Fisher Scientific) according to manufacturer’s protocol. Extracted RNAs were then treated with DNAse (TURBO DNA-free™ kit, Thermo Fisher Scientific) following provider’s instructions. For tumor samples, samples (10–50 mg) were first powdered under liquid nitrogen. For experiments on cell lines, about 3 · 10^5^ cells were used, treatments were performed in triplicates and RNA-seq experiments were performed at least in duplicates. Quality controls were performed using a Bioanalyzer (Agilent) before and after libraries preparation and only RNA samples with RIN (RNA Integrity Number) *>* 6 were used for sequencing. For sequencing library preparation, 1*μg* of highquality total RNA samples were used. Sequencing libraries were prepared using TruSeq Stranded mRNA kit (Illumina) according to manufacturer instructions. Briefly, after purification of poly-A containing mRNA molecules, mRNA molecules were fragmented and reversetranscribed using random primers. dTTP was replaced by dUTP during the second strand synthesis to achieve the strand specificity. Addition of a single A base to the cDNA (dA tailing) was followed by ligation of Illumina adapters. Libraries were quantified by Qubit fluorometer (Life Technologies) and library profiles were assessed using the DNA high Sensitivity LabChip kit on an Agilent Bioanalyzer 2100. Libraries were sequenced on an Illumina Nextseq 500 instrument using 75 base-lengths read chemistry in a paired-end mode. After sequencing, a primary analysis based on AOZAN software64 was applied to demultiplex and control the quality of the raw data. Mapping of the reads to the human reference genome hg38 was performed using STAR v2.5.3a65 with the following parameters: twopassMode, ‘Basic’; chimSegmentMin, 12; chimJunctionOverhangMin, 12; alignSJD-BoverhangMin, 10; alignMatesGapMax, 100000000; alignIntronMax, 200000; alignSJstitchMismatchNmax, 5 -1 5 5.

### RNA extraction and sequencing of cell lines

NCI-H295R (1· 10^6^ cells/well), MUC-1 (5· 10^5^ cells/well) and TVBF-7 (9 · 10^5^ cells/well) cells were plated in 6-well plates (TPP, Trasadin-gen, Switzerland). On the next day, media were replaced with 3ml of the respective media containing forskolin (Sigma-Aldrich) and drugs: 25*μ*M forskolin, 1*μ*M THZ1, 1*μ*M (+)-JQ1, and 40*μ*M mitotane. DMSO concentration in all wells (including controls) was adjusted to 0.5%. Cells were incubated for 24h. RNA was extracted with RNeasy Mini kit (Qiagen, Hilden, Germany) and treated with DNAse (TurboDNA-free™ kit, Thermo Fisher Scientific) using manufacturers protocols. RNA Sequencing was performed by the Genomics Facility Basel, department of Biosystems Science and Engineering (D-BSSE), Basel, Switzerland. Mapping of the reads to the human reference genome hg38 was performed using STAR v2.7.5a with the following parameters: alignIntronMax, 1000000; alignIntronMin, 20: align-MatesGapMax, 1000000: alignSJDBoverhangMin, 1; alignSJover-hangMin, 8; alignSoftClipAtReferenceEnds, Yes; chimJunctionOver-hangMin, 15; chimMainSegmentMultNmax, 1; chimOutType, Junctions SeparateSAMold WithinBAM SoftClip; chimSegmentMin, 15; limitSjdbInsertNsj, 1200000; outFilterIntronMotifs, None; outFilter-MatchNminOverLread, 0.33; outFilterMismatchNmax, 999; outFilterMismatchNoverLmax, 0.1; outFilterMultimapNmax, 20; outFilter-ScoreMinOverLread, 0.33; outFilterType, BySJout; outSAMattributes, NH HI AS nM NM ch; outSAMstrandField, intronMotif; outSAMtype, BAM Unsorted; outSAMunmapped, Withinl; quantMode, TranscriptomeSAM GeneCounts; twopassMode, Basic

### Reduced Representation Bisulfite Sequencing experiments

Genomic DNA was extracted using QIAamp Fast DNA Tissue Kit (Qiagen) according to manufacturer protocol. Reduced Representation Bisulfite Sequencing (RRBS) was performed by Integragen SA (Evry, France) with the Diagenode Premium RRBS kit. In brief, 100 ng of qualified genomic DNA were digested with MspI. After end-repair, A-tailing, and ligation to methylated and indexed adapters, the size selected library fragments were subjected to bisulfite conversion, amplified by PCR, and sequenced on an Illumina NovaSeq sequencer as Paired End 100 bp reads. Image analysis and base calling were performed using Illumina Real Time Analysis (3.4.4) with default parameters. Base calling was performed by using the Real-Time Analysis software sequence pipeline (3.4.4) with default parameters. RRBS data were mapped to the Human genome using BS-Seeker2 (v2.1.8). The following parameters were used: -r (Map reads to the Reduced Representation genome), -c C-CGG (MspI: sites of restriction enzyme and specifying lengths of fragments ranging [40bp, 400bp]. One mismatch was allowed in the adaptor sequence. Allowing local/gapped alignment with Bowtie269 increased the mappability. BS-Seeker2 module bs_seeker2-call_methylation.py was used to call methylation levels from the mapping results with these parameters: rm-SX (Removed reads which would be considered as not fully converted by bisulfite) and rm-overlap (Removed one mate if two mates are overlapped). The methylation callings were saved in the ATCGmap/CGmap format. CGmaptool70 0.1.2 was used to call Differentially Methylated Regions (DMRs).

## References

1. Assié, G. et al. Integrated genomic characterization of adreno-cortical carcinoma. en. Nature Genetics 46, 607–612. ISSN: 1061-4036, 1546-1718 (June 2014).

2. Zheng, S. et al. Comprehensive Pan-Genomic Characterization of Adrenocortical Carcinoma. en. Cancer Cell 29, 723–736. ISSN: 15356108 (May 2016).

3. De Reyniès, A. et al. Gene expression profiling reveals a new classification of adrenocortical tumors and identifies molecular predictors of malignancy and survival. Journal of Clinical Oncology 27. Publisher: J Clin Oncol, 1108–1115. ISSN: 0732183X (Mar. 2009).

4. Giordano, T. J. et al. Molecular classification and prognostication of adrenocortical tumors by transcriptome profiling. Clinical Cancer Research 15. Publisher: American Association for Cancer Research, 668–676. ISSN: 10780432 (Jan. 2009).

5. Hanahan, D. Hallmarks of Cancer: New Dimensions. Cancer Dis-covery 12, 31–46. ISSN: 2159-8274 (Jan. 2022).

6. Barreau, O. et al. Identification of a CpG Island Methylator Phenotype in Adrenocortical Carcinomas. The Journal of Clinical Endocrinology & Metabolism 98. Publisher: Oxford Academic, E174–E184. ISSN: 0021-972X (Jan. 2013).

7. Kerdivel, G., Amrouche, F., Calmejane, M.-A., Hamroune, J. & Boeva, V. DNA hypermethylation driven by DNMT1 and DNMT3A favors tumor immune escape contributing to the aggressiveness of adrenocortical carcinoma. In Revision (Jan. 2023).

8. Hnisz, D. et al. Super-enhancers in the control of cell identity and disease. Cell 155. Publisher: Elsevier B.V., 934. ISSN: 10974172 (Nov. 2013).

9. Boeva, V. et al. Heterogeneity of neuroblastoma cell identity defined by transcriptional circuitries. Nature Genetics 49. Publisher: Nature Publishing Group,1408–1413. ISSN: 15461718 (Sept. 2017).

10. Gartlgruber, M. et al. Super enhancers define regulatory subtypes and cell identity in neuroblastoma. en. Nature Cancer 2, 114–128. ISSN: 2662-1347 (Jan. 2021).

11. Beacon, T. H. et al. The dynamic broad epigenetic (H3K4me3, H3K27ac) domain as a mark of essential genes. Clinical Epigenetics 13, 138. ISSN: 1868-7075 (July 2021).

12. Whyte, W. A. et al. Master transcription factors and mediator establish super-enhancers at key cell identity genes. Cell 153. Publisher: Elsevier B.V., 307–319. ISSN: 10974172 (Apr. 2013).

13. Schnabel, C. A., Selleri, L. & Cleary, M. L. Pbx1 is essential for adrenal development and urogenital differentiation. eng. Gene-sis (New York, N.Y. 37, 123–130. ISSN: 1526-968X (Nov. 2003).

14. Ehrlund, A. et al. Knockdown of SF-1 and RNF31 Affects Components of Steroidogenesis, TGF, and Wnt/-catenin Signaling in Adrenocortical Carcinoma Cells. PLoS ONE 7, e32080. ISSN: 1932-6203 (Mar. 2012).

15. Cerquetti, L. et al. C-MYC modulation induces responsiveness to paclitaxel in adrenocortical cancer cell lines. International Journal of Oncology 46. Publisher: Spandidos Publications, 2231–2240. ISSN: 1019-6439 (May 2015).

16. Dunaief, J. L. et al. The retinoblastoma protein and BRG1 form a complex and cooperate to induce cell cycle arrest. English. Cell 79. Publisher: Elsevier, 119–130. ISSN: 0092-8674, 1097-4172 (Oct. 1994).

17. Peng, L. et al. A Pan-Cancer Analysis of SMARCA4 Alterations in Human Cancers. Frontiers in Immunology 12, 4037. ISSN: 1664-3224 (2021).

18. Passaia, B. d. S. et al. TCF21/POD-1, a Transcritional Regulator of SF-1/NR5A1, as a Potential Prognosis Marker in Adult and Pediatric Adrenocortical Tumors. Frontiers in Endocrinology 9, 38. ISSN: 1664-2392 (2018).

19. Bahr, C. et al. A Myc enhancer cluster regulates normal and leukaemic haematopoietic stem cell hierarchies. en. Nature 553, 515–520. ISSN: 1476-4687 (Jan. 2018).

20. Tibshirani, R. The lasso method for variable selection in the Cox model. Statistics in medicine 16. Publisher: Wiley Online Library, 385–395 (1997).

21. Lee, D. D. & Seung, H. S. Learning the parts of objects by non-negative matrix factorization. en. Nature 401, 788–791. ISSN: 1476-4687 (Oct. 1999).

22. Robinson, M. D., McCarthy, D. J. & Smyth, G. K. : a Bioconductor package for differential expression analysis of digital gene expression data. Bioinformatics 26, 139–140. ISSN: 1367-4803 (Jan. 2010).

23. Saint-André, V. et al. Models of human core transcriptional regulatory circuitries. Genome Research 26. Publisher: Cold Spring Harbor Laboratory Press, 385–396. ISSN: 15495469 (Mar. 2016).

24. Lin, C. Y. et al. Active medulloblastoma enhancers reveal subgroup-specific cellular origins. Nature 530. Publisher: Nature Publishing Group, 57–62. ISSN: 14764687 (Feb. 2016).

25. Ashoor, H. et al. HMCan: a method for detecting chromatin modifications in cancer samples using ChIP-seq data. Bioinformatics 29. Publisher: Oxford Academic, 2979–2986. ISSN: 1460-2059 (Dec. 2013).

26. The FANTOM Consortium et al. An atlas of active enhancers across human cell types and tissues. en. Nature 507, 455–461. ISSN: 0028-0836, 1476-4687 (Mar. 2014).

27. Gazdar, A. F. et al. Establishment and Characterization of a Human Adrenocortical Carcinoma Cell Line That Expresses Multiple Pathways of Steroid Biosynthesis1. Cancer Research 50, 5488–5496. ISSN: 0008-5472 (Sept. 1990).

28. Hantel, C. et al. Targeting heterogeneity of adrenocortical carcinoma: Evaluation and extension of preclinical tumor models to improve clinical translation. en. Oncotarget 7. Publisher: Impact Journals, 79292–79304. ISSN: 1949-2553 (Oct. 2016).

29. Sigala, S. et al. A Comprehensive Investigation of Steroidogenic Signaling in Classical and New Experimental Cell Models of Adrenocortical Carcinoma. en. Cells 11. Number: 9 Publisher: Multidisciplinary Digital Publishing Institute, 1439. ISSN: 2073-4409 (Jan. 2022).

30. Wu, T., Huang, H. & Wang, X. Dissecting super-enhancer heterogeneity: time to re-examine cancer subtypes? English. Trends in Genetics 38. Publisher: Elsevier, 1199–1203. ISSN: 0168-9525 (Dec. 2022).

31. Crona, J. & Beuschlein, F. Adrenocortical carcinoma — towards genomics guided clinical care. Nature Reviews Endocrinology 15. Publisher: Nature Publishing Group, 548–560. ISSN: 17595037 (Sept. 2019).

32. Cai, Y. et al. H3K27me3-rich genomic regions can function as silencers to repress gene expression via chromatin interactions. en. Nature Communications 12, 719. ISSN: 2041-1723 (Jan. 2021).

33. Zhang, T., Cooper, S. & Brockdorff, N. The interplay of histone modifications – writers that read. EMBO Reports 16, 1467–1481. ISSN: 1469-221X (Nov. 2015).

34. Drelon, C. et al. EZH2 is overexpressed in adrenocortical carcinoma and is associated with disease progression. Human Molecular Genetics 25, 2789–2800. ISSN: 0964-6906 (July 2016).

35. Zhou, V. W., Goren, A. & Bernstein, B. E. Charting histone modifications and the functional organization of mammalian genomes. en. Nature Reviews Genetics 12, 7–18. ISSN: 1471-0056, 1471-0064 (Jan. 2011).

36. Sengupta, S. & George, R. E. Super-Enhancer-Driven Transcriptional Dependencies in Cancer. en. Trends in Cancer 3, 269–281. ISSN: 2405-8033 (Apr. 2017).

37. Mohan, D. R. et al. Beta-catenin programs a tissue-specific epigenetic vulnerability in aggressive adrenocortical carcinoma en. Pages: 2022.07.02.497654 Section: New Results. July 2022.

38. Dong, C.-H. et al. LMNB2 promotes the progression of colorectal cancer by silencing p21 expression. en. Cell Death & Disease 12, 1–12. ISSN: 2041-4889 (Mar. 2021).

39. Bothou, C. et al. Novel Insights into the Molecular Regulation of Ribonucleotide Reductase in Adrenocortical Carcinoma Treatment. en. Cancers 13. Number: 16 Publisher: Multidisciplinary Digital Publishing Institute, 4200. ISSN: 2072-6694 (Jan. 2021).

40. Sigala, S. et al. An update on adrenocortical cell lines of human origin. Endocrine 77, 432–437. ISSN: 1355-008X (2022).

41. Creemers, S. G. et al. Inhibition of Human Adrenocortical Cancer Cell Growth by Temozolomide in Vitro and the Role of the MGMT Gene. en. The Journal of Clinical Endocrinology & Metabolism 101, 4574–4584. ISSN: 0021-972X, 1945-7197 (Dec. 2016).

42. Gerson, S. L. Clinical Relevance of MGMT in the Treatment of Cancer. Journal of Clinical Oncology 20. Publisher: Wolters Kluwer, 2388–2399. ISSN: 0732-183X (May 2002).

43. Weiss, L. M. Comparative histologic study of 43 metastasizing and nonmetastasizing adrenocortical tumors. eng. The American Journal of Surgical Pathology 8, 163–169. ISSN: 0147-5185 (Mar. 1984).

44. Stell, A. & Sinnott, R. The ENSAT registry: a digital repository supporting adrenal cancer research. Studies in health technology and informatics 178, 207–12 (July 2012).

45. R Core Team. R: The R Project for Statistical Computing Vienna, Austria, 2021.

46. Benjamini, Y. & Hochberg, Y. Controlling the False Discovery Rate: A Practical and Powerful Approach to Multiple Testing. Journal of the Royal Statistical Society: Series B (Methodological) 57. Publisher: Wiley, 289–300. ISSN: 2517-6161 (Jan. 1995).

47. Boeva, V. et al. Control-FREEC: a tool for assessing copy number and allelic content using next-generation sequencing data. Bioinformatics 28. Publisher: Oxford Academic, 423–425. ISSN: 1460-2059 (Feb. 2012).

48. Amemiya, H. M., Kundaje, A. & Boyle, A. P. The ENCODE Blacklist: Identification of Problematic Regions of the Genome. Scientific Reports 9. Publisher: Nature Publishing Group, 1–5. ISSN: 20452322 (Dec. 2019).

49. Polit, L. et al. CHIPIN: ChIP-seq inter-sample normalization based on signal invariance across transcriptionally constant genes. BMC Bioinformatics 22, 407. ISSN: 1471-2105 (Aug. 2021).

50. Howe, K. L. et al. Ensembl 2021. en. Nucleic Acids Research 49, D884–D891. ISSN: 0305-1048, 1362-4962 (Jan. 2021).

51. Rao, S. S. et al. A 3D map of the human genome at kilobase resolution reveals principles of chromatin looping. Cell 159. Publisher: Cell Press, 1665–1680. ISSN: 10974172 (Dec. 2014).

52. Lambert, S. A. et al. The Human Transcription Factors. Cell 172. Publisher: Cell Press, 650–665. ISSN: 10974172 (Feb. 2018).

53. Forbes, S. A. et al. COSMIC: Somatic cancer genetics at highresolution. Nucleic Acids Research 45. Publisher: Oxford University Press, D777–D783. ISSN: 13624962 (Jan. 2017).

54. Quintero, A. et al. ShinyButchR: Interactive NMF-based decomposition workflow of genome-scale datasets. Biology Methods & Protocols 5, bpaa022. ISSN: 2396-8923 (Oct. 2020).

55. Cox, D. R. Regression models and life-tables. Journal of the Royal Statistical Society: Series B (Methodological) 34. Publisher: Wiley Online Library, 187–202 (1972).

56. Meinshausen, N. & Bühlmann, P. Stability selection. Journal of the Royal Statistical Society: Series B (Statistical Methodology) 72. Publisher: Wiley Online Library, 417–473 (2010).

